# Single oocyte full-length isoform sequencing unveils the impact of transposable elements on RNA diversity and stability during oocyte maturation

**DOI:** 10.1101/2025.06.17.659919

**Authors:** Yuqian Wang, Wei Wang, Yujun Liu, Yiming He, Hongyu Song, Ming Yang, Nan Wang, Xiaomeng Wang, Ling Ding, Ying Kuo, Yuwen Xiu, Zhengrong Du, Lu Chen, Ying Lian, Qiang Liu, Liying Yan, Jie Qiao, Peng Yuan

**Author notes:** Correspondence (P.Y.); (J.Q.); (L.Y.); (Q.L.). These authors contributed equally (Yuqian Wang, Wei Wang, Yujun Liu, Yiming He, Hongyu Song).

## Abstract

The oocyte-specific isoforms play crucial roles in oocyte maturation, while current understanding of the oocyte transcriptome is mainly focused on gene level. Here, we utilized single cell full-length isoform sequencing based on third generation sequencing to detect entire transcripts in human and mouse oocytes. Isoform diversity during oocyte maturation was systematically profiled, including 6,736 and 4,902 putative novel human and mouse transcripts, respectively. More than half of novel isoforms were categorized as the novel-not-in-catalog (NNC) and may serve specific functions in oocytes, including novel isoforms of *ARHGAP18*, colocalized with microtubules and targeted knockdown disrupting oocyte maturation. Moreover, over 25% of NNC isoforms were derived from transposable elements (TEs), and their incorporation within transcripts could enhance isoform stability during oocyte maturation. Altogether, our findings represent a valuable resource showcasing the complexity and diversity of RNA isoforms in oocytes, as well as TE co-option for novel isoform generation and isoform stability enhancement.

## Introduction

The oocyte is the largest cell type in most organisms, with abundant RNAs synthesized and accumulated in cytoplasm to support oocyte maturation and early embryo development (Cheng et al., 2022; Su et al., 2007; Wan et al., 2023). Upon initiation of oocyte maturation, the germinal vesicle (GV) stage oocyte (arrested in prophase I of meiosis), undergoes the germinal vesicle break-down (GVBD) and then resumes meiosis I, leading to the intermediate maturation stage metaphase I (MI). As maturation proceeds further, a metaphase II (MII) stage oocyte with abundant mRNAs and a small first polar body are produced through asymmetric cell division (Do et al., 2018; Kocabas et al., 2006). Given the pivotal role of RNA in regulating oocyte maturation (Llonch et al., 2021; Pietroforte et al., 2023), investigating the diversity of RNA expression and regulation underlying oocyte maturation is crucial.

Previous studies provided valuable insight into gene-level abundance, complexity and dynamics of transcriptome during oocyte maturation through next generation sequencing (NGS) (Hu et al., 2022; Llonch et al., 2021; Zhang et al., 2018). Although it is transcriptional silencing after oocyte meiotic resumption, selective isoform usage and oocyte specific isoforms are critical for regulating the oocyte maturation (He et al., 2021; Pietroforte et al., 2023). However, the isoform-level diversity and isoform usage characteristics have not been explored in detail. Further studies at the isoform level are imperative, given its richness in terms of RNA types and content. The transcriptome complexity of oocyte maturation primarily results from alternative splicing (AS), leading to the generation of oocyte-specific transcripts (Do et al., 2018). Some oocyte-specific transcripts have been identified and confirmed to have crucial functions, such as ELAVL2^O^ is translated by an oocyte-specific transcript, which functions as a repressor of translation that contributes to the acquisition of developmental competence of oocytes (Chalupnikova et al., 2014). As another example, long MEX3C isoform is specifically expressed in mouse oocytes and have been shown to enhance the nuclear export and mRNA stability of *FOS* so as to preserve maternal mRNA for subsequent translation in early embryos (Li et al., 2016). Above functional studies demonstrate the functional importance of oocyte-specific transcripts during oocyte maturation, and motivate a systematic exploration of the diversity of oocyte-specific transcripts.

Long dismissed as ‘junk DNA’, transposable elements (TEs), which make up approximately 40% of the genomes of human and mouse (Miao et al., 2020), are recognized as functional cis-regulatory elements in early mammalian development (Do et al., 2018; Fueyo et al., 2022; Georgiou et al., 2009). The incorporation of TE sequences within transcripts has proven to be a considerable source of germline-specific isoforms and is crucial for oocyte maturation and early embryonic development (Franke et al., 2017; Modzelewski et al., 2021; Peaston et al., 2004). Illustratively, an MT-C retrotransposon could serve as alternative promoter to generate an N-terminally truncated Dicer^O^ isoform in mouse oocytes, endowing the oocyte-specific isoform of *Dicer* with more efficient endonucleolytic cleavage activity compared to its counterparts in somatic cells (Flemr et al., 2013). Previous studies on the contribution of TE sequences to the transcriptome have been challenging due to the repetitive nature of TEs and short read lengths of NGS technique (Modzelewski et al., 2021; Treangen and Salzberg, 2011). Thus, an approach to the comprehensive profiling of TE-derived isoforms during oocyte maturation is urgently needed.

Full-length isoform sequencing through third generation sequencing (TGS) platform has been a key approach to overcome existing limitations inherent to NGS approach that affected read length (Amarasinghe et al., 2020; Uapinyoying et al., 2020). Indeed, TGS approach enables the detection of full-length transcripts, and can also map the incorporation of TE sequences within transcripts (Berrens et al., 2022). Novel isoforms and AS events in multiple tissues have been characterized (Fan et al., 2020; Glinos et al., 2022; Qiao et al., 2020; Sun et al., 2021), and recent research has also unveiled the complexity of transcriptome in human embryos (Torre et al., 2023). Due to the scarcity of human oocyte samples and the short-read limitation of NGS, the overall profiling of oocyte-specific transcripts, as well as the comparative analyses between human and mouse oocytes has likely been underestimated. In this study, we performed a single-cell full-length transcript profiling study to systematically characterize RNA isoforms and the incorporation of TE sequences within transcripts during human and mouse oocyte maturation.

## Results

### 1. Comprehensive sequencing of full-length RNA isoforms in human and mouse oocytes

We implemented PacBio long-read isoform sequencing on human and mouse oocytes to identify full-length isoforms. An overview of sample collection and sequencing methodology is shown in Fig. 1A. Human oocytes at three representative stages (GV, MI, MII) and mouse oocytes at four representative stages (GV, GVBD, MI, MII) were collected. After reverse transcription in which MMLV reverse transcriptase was employed, the 5’ capped mature transcripts were highly preferentially combined and reversed. Barcodes and unique molecular identifiers (UMIs) were introduced to label each individual oocyte prior to polymerase chain reaction (PCR) amplification. Labeled cDNA products from a total of 20 human oocytes and 26 mouse oocytes were pooled and PacBio adaptors were ligated to the cDNA products for isoform sequencing. Through this approach, a total of approximately 6 Gb of raw data was generated and 15 million circular consensus sequence (CCS) reads were obtained (Supplementary Fig. 1A,B). More than 90 percent of demultiplex reads contain both 5’ PCR oligo and 3’ poly A tail (Supplementary Fig. 1C). A total of 11.0 million qualified and deduplicated reads were obtained (Supplementary Fig. 1B) and the mean length of these high-quality reads was approximately 1800 bp, and nearly half of these reads reached Q50 (Supplementary Fig. 1D). The depth, accuracy, reproducibility and the uniformity of reads coverage of this sequencing were assessed to be sufficient for downstream analysis (Supplementary Fig. 1C-E). Next, we systematically analyzed the differences between NGS and TGS data at the isoform level. The quality of transcripts was assessed based on transcript length, exon count and transcript completeness (CAGE and Poly A supported transcripts), and the results demonstrated that the quality of transcripts obtained by TGS was distinctly improved over NGS and GENCODE assemblies (Fig. 1B and Supplementary Fig. 1F). Altogether, these results demonstrate the high throughput, sensitivity, and integrity of the characterization and quantification of full-length isoform sequencing from single oocyte.

**Figure 1.**
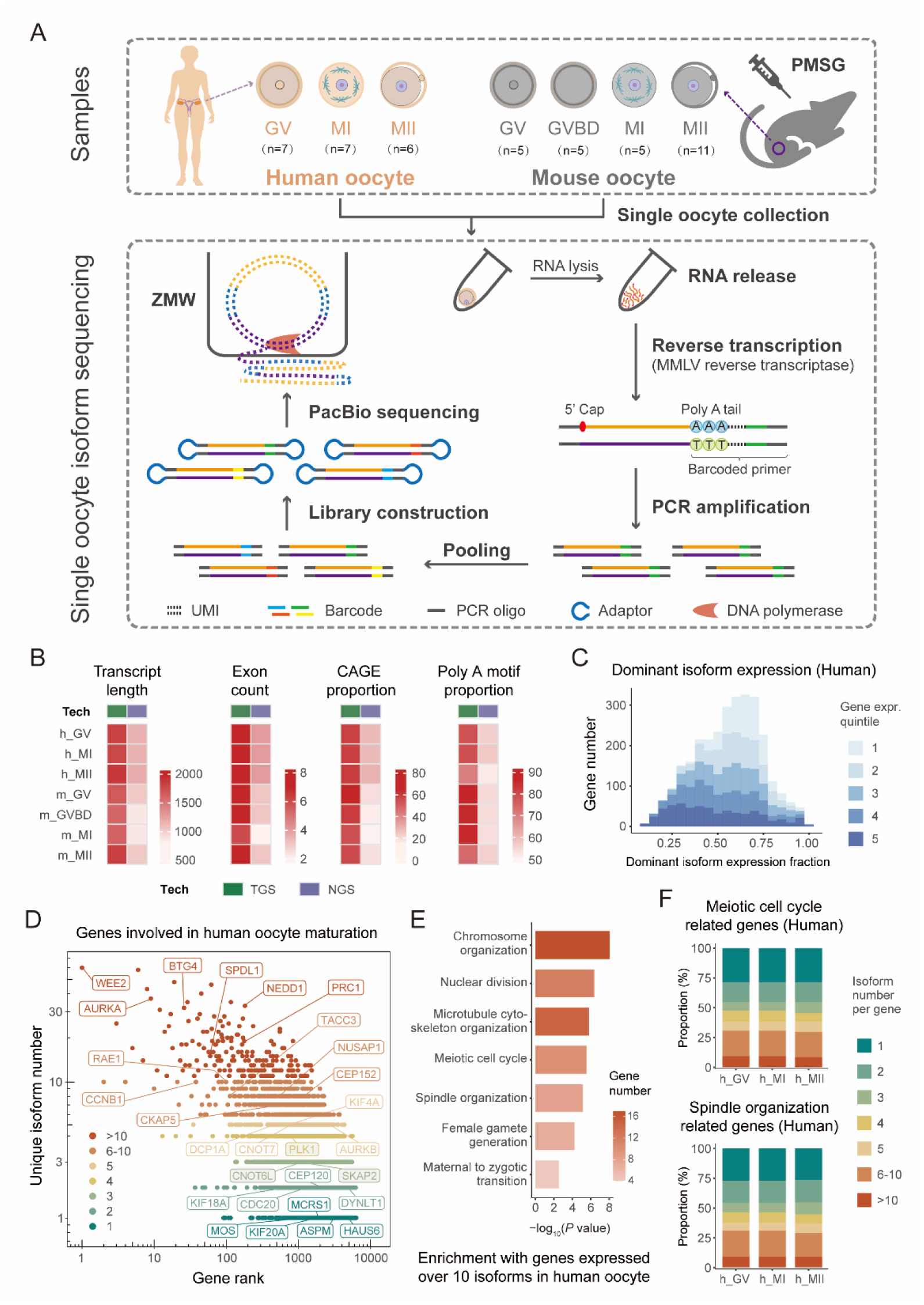
Methodology overview of full-length isoform sequencing and basic analysis of expression in human and mouse oocytes. (A) Schematic diagram of sample collection, full-length isoform library construction, and sequencing. PMSG, pregnant mare serum gonadotropin; ZMW, zero-mode waveguide. (B) Pileup heatmaps showing the comparison between transcriptome assembled using TGS and NGS, including median transcript length (excluding introns), median exon number, CAGE and poly A support for isoforms. (C) Dominant isoform expression fraction of multi-isoform genes stratified by gene expression percentiles. The highest expressed isoform of a certain gene was defined as dominant isoform (D) Unique isoform number detected of all genes ranked by gene expression in human oocyte. Genes involved in oocyte maturation were labeled. (E) Representative pathway enrichment of genes with more than 10 unique isoforms in human oocyte. (F) Isoform diversity of genes related to meiotic cell cycle and spindle organization detected during human oocyte maturation.

To ensure the reliability and accuracy of the transcripts detected in oocytes noted for further study, we only included the isoforms that were present in at least three samples, as previously reported (Fan et al., 2020), and this approach yielded a total of 16,769 isoforms in human oocytes and 14,987 in mouse oocytes. The enrichment of CAGE and polyA-seq signals at the assembled transcripts demonstrates the robustness of our assemblies (Supplementary Fig. 1G). Given the high-quality and reliable isoforms through our selection criteria, we sought to characterize the isoform diversity and their usage patterns during oocyte maturation. The correlation analysis on isoform expression level showed that human oocytes at MII stage are significantly different from oocytes at GV and MI stages (Supplementary Fig. 1H). While, during mouse oocyte maturation, oocytes at GV, GVBD-MI, and MII stages are distinguishable at the isoform level (Supplementary Fig. 1H). For multi-isoform genes in human oocytes, we quantified the distribution of dominant isoform expression fractions, revealing that the dominant isoforms exhibited a relatively even pattern of expression despite the gene expression levels, and this indicates the complexity of isoform usage patterns during oocyte maturation (Fig. 1C). Additionally, we calculated the number of detected isoforms for each gene and ranked genes according to their expression levels (Fig. 1D). Notably, many genes involved in oocyte maturation were found to exhibit high isoform diversity, such as *WEE2* and *AURKA,* which are crucial for oocyte meiosis process (Sang et al., 2018; Schindler et al., 2012). Further enrichment analysis of genes with over 10 distinct isoforms revealed significant enrichment in pathways essential for oocyte maturation, such as chromosome organization, meiotic cell cycle, and spindle organization (Fig. 1E). Among the meiotic cell cycle and spindle organization related genes we detected in human oocytes, almost half of these genes expressed more than 3 isoforms, and a quarter expressed over 5 isoforms (Fig. 1F), contrasting with that only 5-11% genes expressed more than 5 isoforms in human granulosa cells (GCs) and embryonic stem cells (ESCs) (Supplementary Fig. 1I). These patterns indicated high isoform diversity in genes related to meiotic cell cycle and spindle organization in oocytes, which remains consistent across the GV, MI, and MII stages during oocyte maturation (Fig. 1F). Altogether, these results demonstrate that our long-read sequencing approach provides comprehensive insights into isoform level dynamics underlying the physiological processes of oocyte maturation.

### 2. Full-length isoform sequencing identifies the overall pattern of human oocytes and reveals its diversity

We compared our data with previous studies and GENCODE database (Franke et al., 2017; Hu et al., 2022; Liu et al., 2019; Llonch et al., 2021; Zhang et al., 2018; Zhao et al., 2020), and found that over 36% of the isoforms we identified were not detected in these datasets (Supplementary Fig. 2A). Next, we used SQANTI3 to further annotate and classify these isoforms (Fig. 2A). The results demonstrated that 59.8% of detected transcripts in human oocytes were identified as full splice match (FSM) transcripts with exact matches to all splice junctions, or incomplete splice match (ISM) transcripts that matched the reference splice junctions partially (Fig. 2B and Supplementary Table 1). The distribution of known transcripts (FSM and ISM) was found to be largely similar among the three human oocyte stages (Fig. 2B). We also detected novel isoforms, characterized as novel in catalog (NIC), novel not in catalog (NNC), fusion, genic genomic categories, as well as antisense and intergenic categories that may contribute to novel genes (Fig. 2A,B). In total, 6,736 novel transcripts were identified, and comprised 40.2% of all transcripts in human oocytes (Supplementary Table 1), whereas novel isoforms only occupied around 10-12% in human GCs and ESCs (Supplementary Fig. 2B and Supplementary Table 1). After evaluating the coding probability of the detected isoforms (Fig. 2C), we further analyzed the proportions of coding and non-coding transcripts within each isoform category. The FSM, ISM, NNC and NIC categories were predominantly composed of coding isoforms (accounting for over 70%). In contrast, the antisense and intergenic categories were largely composed of non-coding transcripts (greater than 88.9%) (Supplementary Fig. 2C and Supplementary Table 2). Furthermore, the distribution of coding and non-coding transcripts was observed to be relatively consistent across the three stages of human oocyte (Supplementary Fig. 2C).

**Figure 2.**
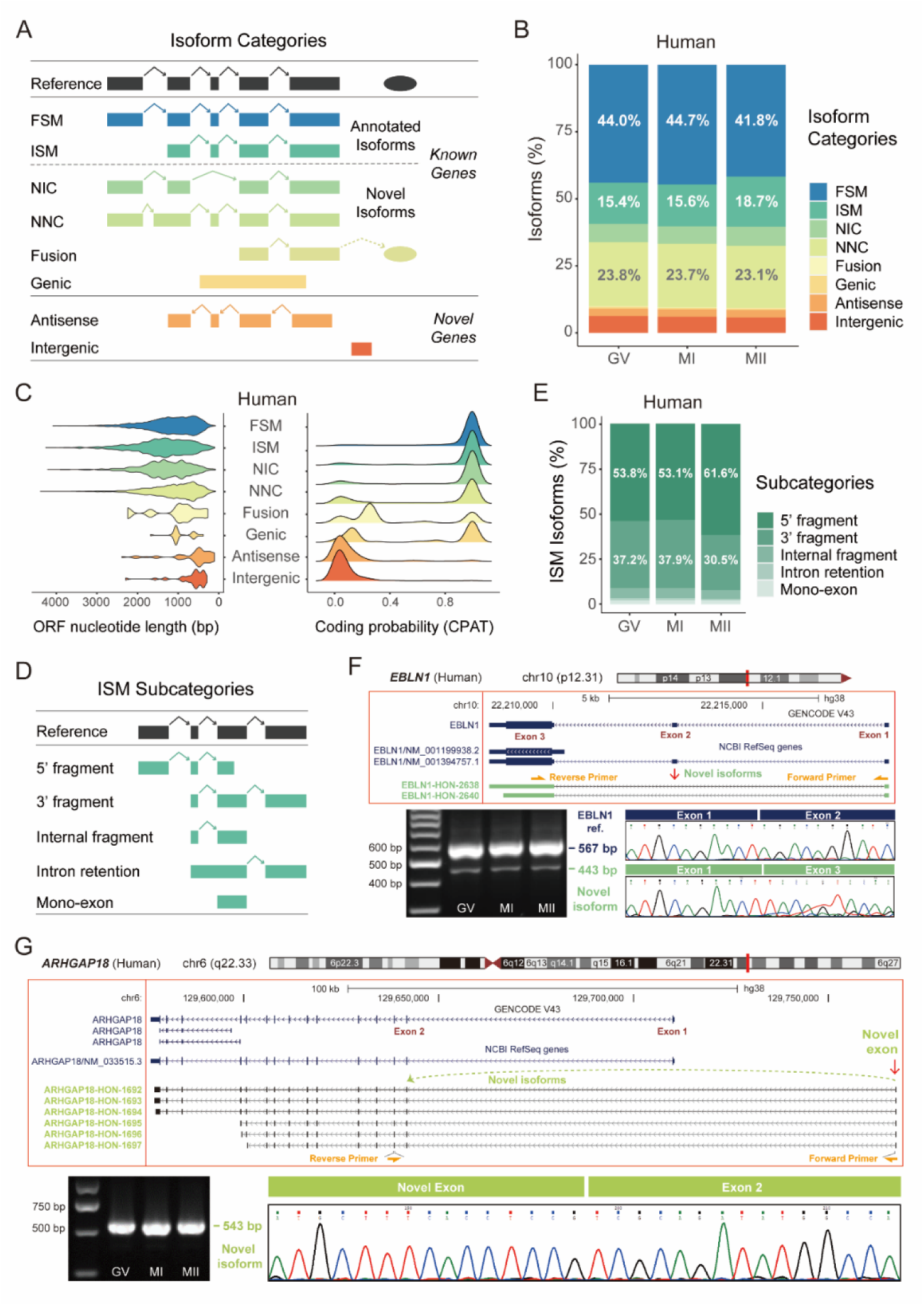
Isoform categories profiling and validation of novel isoforms in human oocytes. (A) Schematic diagram of isoform structural categories. FSM, full splice match; ISM, incomplete splice match; NIC, novel in catalog; NNC, novel not in catalog. (B) Isoform percentages of different categories of all detected isoforms during human oocyte maturation. Each isoform has a minimum of three biological replicates. (C) Distribution of open reading frame (ORF) length and coding probability of isoforms by category. (D) Schematic diagram of ISM subcategories. (E) Percentages of ISM subcategories in different maturation stages of human oocytes. (F) The location in the genome and the isoform structure of *EBLN1* are shown in the upper part. The red arrow indicates the absent exon of the novel isoforms, and the orange arrows showed the position of primers used for PCR. The lower left shows the validation of *EBLN1* novel isoforms in different stages of development in human oocytes by agarose gel electrophoresis after PCR. Sanger sequencing chromatogram (lower right) confirming the splice junctions of PCR products. (G) The location in the genome and the isoform structure of *ARHGAP18* are shown in the upper part and the red arrows show the additional novel exons. The orange arrows showed the position of primers used for PCR. The lower left displays the *ARHGAP18* novel isoforms in different stages of development in human oocytes using agarose gel electrophoresis following PCR. The lower right shows splice junction between novel exon and exon 2.

From our analysis using SQANTI3, FSM and ISM transcripts could each be characterized as five subcategories (Fig. 2D and Supplementary Fig. 2D). Among FSM, five subcategories remained stable in human oocyte maturation (Supplementary Fig. 2E). Among ISM, the 5 prime fragment subcategory was found to encompass more than half across all human oocyte stages and its proportion increased from the GV to the MII stages, while the 3 prime fragment subcategory correspondingly decreased, suggesting a switch from 3 prime fragment to 5 prime fragment isoforms during human oocyte maturation (Fig. 2E). To confirm the authenticity of the ISM category, a 3 prime fragment ISM isoform of *ELOVL5* gene was validated through PCR using specific targeted primers, followed by Sanger sequencing (Supplementary Fig. 2F). The results showed that the 3 prime fragment ISM isoform of *ELOVL5*, which features an alternative 5 prime end, representing it a complete isoform in oocytes rather than a degradation artifact (Supplementary Fig. 2F). In terms of novel isoforms, NNC (23.6% of all isoforms in human oocytes) was a dominant category for transcripts that had at least one donor or acceptor site that was not annotated (Fig. 2A, Supplementary Fig. 2G and Supplementary Table 1), while the NNC category in human GCs and ESCs only accounted for 3% of all transcripts detected (Supplementary Fig. 2B). The high proportion of novel isoforms classified as NNC in human oocytes (58.7%) suggested that such novel sequences may have hitherto unknown functions during oocyte maturation (Fig. 2B, Supplementary Fig. 2H and Supplementary Table 1). The increased ISM proportion in known isoforms and the preponderance of NNC category in novel isoforms suggest that isoform composition is diversiform and unique during oocyte maturation.

To verify the robustness of our technique, two novel isoforms were selected for validation. As shown in Fig. 2F, novel transcripts of the *EBLN1* gene characterized as NIC, were validated through PCR, followed by agarose gel electrophoresis and Sanger sequencing. Compared with the known transcript, the novel isoform exhibited alternative splicing that excluded exon 2, and this novel isoform was indeed expressed in human oocytes (Fig. 2F). We also validated the novel isoforms of *ARHGAP18* gene identified as NNC. As shown, Sanger sequencing demonstrated that the novel exon was attached to known exon 2, excluded known transcript exon 1 and included a new exon located approximately 50Kb upstream of the exon 1 (Fig. 2G). The agarose gel electrophoresis result showed that the novel isoform was authentically expressed in human oocytes (Fig. 2G). The oocyte-specific expression of novel isoforms, especially the NNC transcripts with novel sequences, suggests their potential roles in oocyte maturation.

### 3. The categories distribution and diversity between human and mouse oocytes are conservative

Using the same analysis pipeline with human oocytes (Supplementary Fig. 1A), we analyzed our data from single mouse oocytes and discovered a total of 4,902 novel transcripts (Supplementary Table 1), isoform categories across four developmental stages (GV, GVBD, MI, MII), as shown in Fig. 3A. The annotated isoforms, including FSM and ISM, account for 67.3% of all isoforms in mouse oocytes (Fig. 3A). Compared to human oocytes, the relative proportions of isoform categories were largely of similar distribution, while the proportion categorized as FSM was found to be higher in mouse versus human oocyte (Fig. 3A, B). When we looked for variations in isoform categories during oocyte maturation, we found a category switch from FSM to ISM across oocyte maturation in both human and mouse (Fig. 3C). Analysis of subcategories of FSM and ISM showed these to be relatively unchanged during oocyte maturation in mouse (Supplementary Fig. 3A,B). Notably, the 3 prime fragment subcategory constitutes the main component of ISM in mouse oocytes, while the 5 prime fragment subcategory constitutes the majority of ISM in human oocytes (Fig. 3D and Supplementary Fig. 3C). With regards to novel isoforms, a significant feature of both mouse and human oocytes was that the NNC category occupied the majority (Fig. 3B). Moreover, more than 95% of NNC transcripts contained at least one novel splice site rather than intron retention, which reflects the authenticity and accuracy of this method (Supplementary Fig. 3D). These findings indicate that the spectrum of isoforms shows great similarity between human and mouse oocytes.

**Figure 3.**
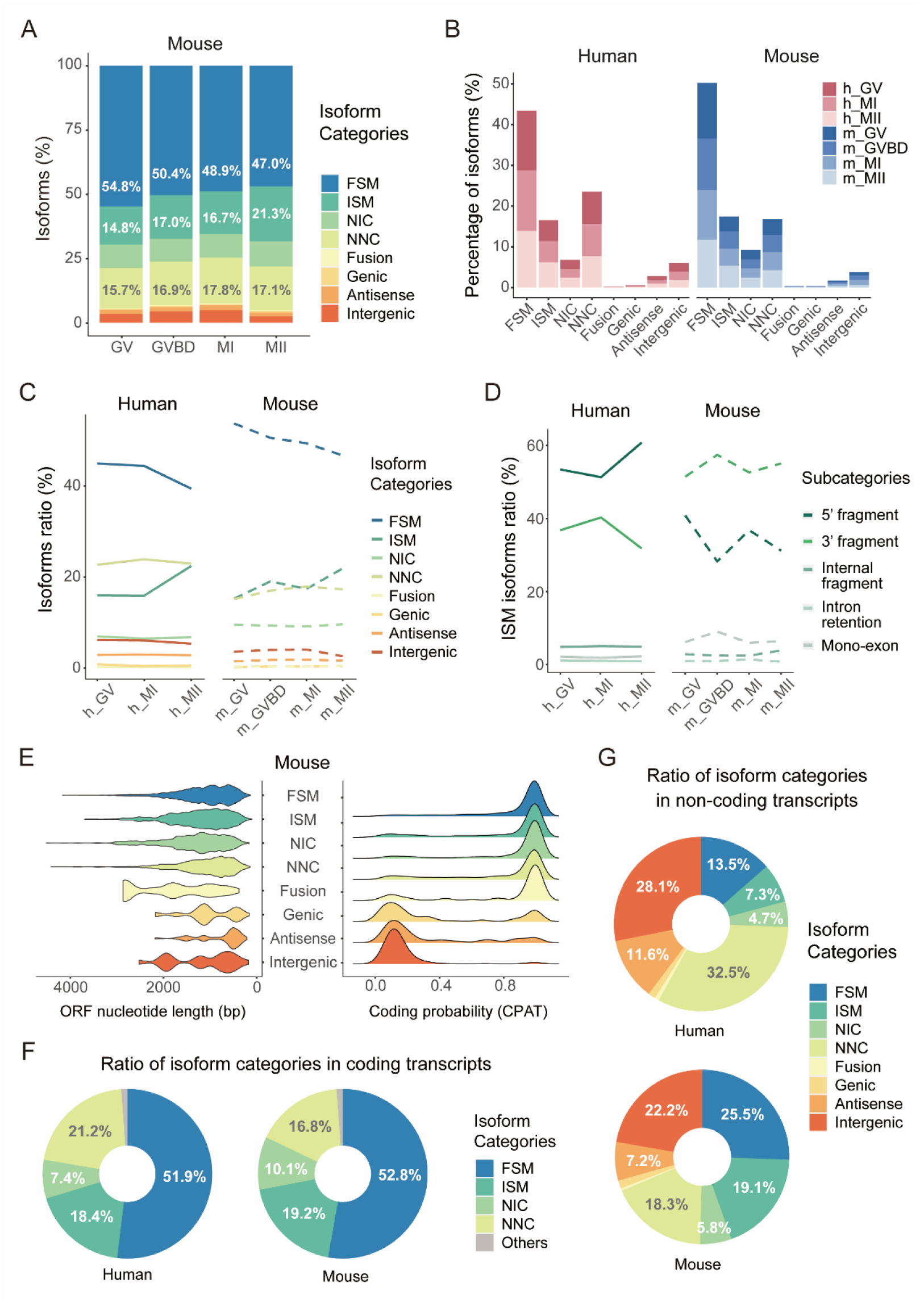
Profiling of different structures of isoforms in human oocytes and comparison between human and mouse oocytes. (A) Percentages of different categories of all detected isoforms during mouse oocyte maturation. Each isoform has a minimum of three biological replicates. (B) Percentages of each isoform category in three human oocyte maturation stages (left) and four mouse oocyte maturation stages (right). (C) Line chart showing the median ratios of different category dynamics during human and mouse oocyte maturation. (D) Line chart showing the median ratios of different ISM subcategory dynamics during human and mouse oocyte maturation. (E) Distribution of mouse ORF length and coding probability of isoforms by category. (F,G) Distribution of each isoform categories in coding (F) and non-coding (G) transcripts of human and mouse oocytes.

In addition, the isoform coding probabilities and distribution of coding and non-coding transcripts were largely similar between human and mouse oocytes (Fig. 3E, Supplementary Fig. 2C and Supplementary Fig. 3E). In both species, the FSM, ISM, NIC and NNC categories were predominantly composed of coding isoforms (Supplementary Fig. 2C and Supplementary Fig. 3E and Supplementary Table 2). Further analysis of isoform category distribution in coding and non-coding transcripts revealed that FSM, ISM, NIC and NNC accounted for 98.9% of all coding transcripts in both human and mouse oocytes (Fig. 3F). The proportion of genic, antisense and intergenic categories was higher in non-coding transcripts compared to their minimal proportions in coding transcripts across both species (Fig. 3G). This isoform analysis therefore suggests that the category, distribution and diversity of transcripts between human and mouse oocytes are similar and conservative. Furthermore, the ISM of known transcripts and the NNC of novel transcripts are the dominant categories, which is a significant signature in both human and mouse oocytes.

### 4. Identifying the DTU events during human and mouse oocyte maturation

Previous studies focused on analyzing the diversity of human and mouse at the gene level based on NGS, and may have overlooked the isoform characteristics in oocyte maturation (Llonch et al., 2021; Zhang et al., 2018). Here, we employed full-length isoform sequencing to comprehensively profile the isoform expression in human and mouse oocyte, and we found the total mRNA showing a degradation trend during oocyte maturation (Supplementary Fig. 4A). Particularly, mRNAs significantly decreased at the MII stage, prompting further comparison of transcriptional dynamics between MII and earlier stages. The differential analyses between GV-MI and MII stages at isoform and gene level indicate some differences and isoform level comparison can provide higher-dimensional information (Fig. 4A, Supplementary Fig. 4B). At the isoform level, we analyzed the isoform changes between the GV-MI and MII stages, and identified 720 isoforms with significant fraction changes corresponding to 398 differential transcript usage (DTU) host genes in human, and 649 isoforms with changes corresponding to 344 DTU host genes in mouse. Analysis of DTU-associated isoforms identified that approximately 50% in human and 40% in mouse originated from non-canonical transcripts, predominantly categorized as NNC and ISM types (Fig. 4B,C). A subset of DTU host genes in both species enriched in pathways essential for oocyte maturation, including cell cycle and RNA metabolism (Supplementary Fig. 4C). We found 22 overlapped DTU host genes between human and mouse (Fig. 4D). Notably, eIF3e, a subunit of the mammalian eukaryotic translation initiation factor 3 complex essential for early embryonic development and cell proliferation (Sadato et al., 2018), showed preferential isoforms usage in both human and mouse (Supplementary Fig. 4D). During oocyte maturation, almost all differentially expressed genes (DEGs) exhibited a trend towards degradation, however, some DTU events were observed at the MII stage with increased isoform fraction at the transcript level (Supplementary Fig. 4E).

**Figure 4.**
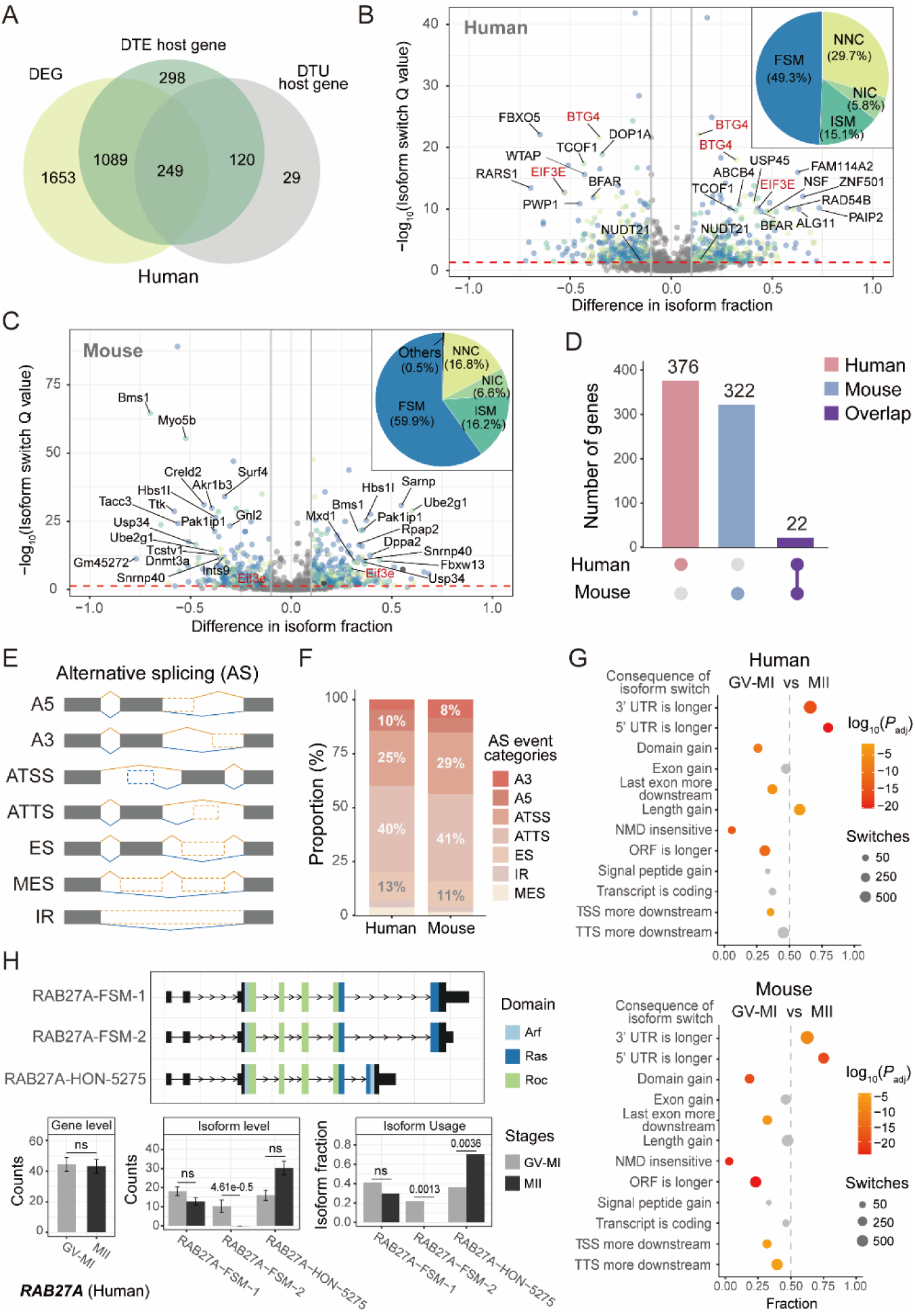
Isoform switching dynamics during oocyte maturation in human and mouse. (A) Venn diagram illustrating the overlap of genes with significant DEG, DTE and DTU during human oocyte maturation. (B,C) Volcano plot depicting isoform switching during human (B) and mouse (C) oocyte maturation respectively. Dot colors represent isoform categories, as indicated in the pie chart (upper right), with non-significant isoforms shown in gray. The pie chart displays the percentage of isoform categories among all detected DTUs. (D) UpSet plot highlighting the overlap of DTU host genes between human and mouse oocytes. (E) Schematic diagram illustrating various categories of alternative splicing (AS) events. (F) The proportion of different types of AS events in human and mouse DTUs. (G) Consequence enrichment of isoform switches between GV-MI and MII stages in human (upper) and mouse (lower). *P* values from two-sided binomial test with FDR-adjusted are highlighted from orange to red, while non-significant changes are shown in grey. The dot size indicates the number of isoforms associated with each type of consequence. (H) An example of DTU events observed within gene *RAB27A* in human oocyte. Isoform structures of *RAB27A* (upper), gene and isoform level expression (lower left and middle), and isoform fraction (lower right) are shown. DEG, differentially expressed gene; DTE, differential transcript expression; DTU, differential transcript usage; A5, alternative 5’ donor site; A3, alternative 3’ acceptor site; ATSS, alternative transcription start site; ATTS, alternative transcription termination site; ES, exon skipping; MES, multiple exon skipping; IR, intron retention.

AS serves as an important mechanism engendering isoform diversity (Baralle and Giudice, 2017). According to the categories of AS (Fig. 4E), we analyzed the isoforms involved in DTU events of human and mouse, suggesting that the AS events of DTU isoforms are similar across the two species. Alternative transcription termination sites (ATTS) and alternative transcription start sites are the most frequent AS events during oocyte maturation in both species, comprising over 65% of all detected AS events (Fig. 4F). In both human and mouse, a large number of ATTS gain events were documented during oocyte maturation, along with the occurrence of elongated untranslated regions (UTRs) at the MII stage (Supplementary Fig. 4F, Fig. 4G). For instance, RAB27A is the member of the Rab family of vesicular trafficking proteins that has been reported to regulate cortical granule exocytosis after MII oocyte activation in mouse (Wang et al., 2016). We found that, while the overall expression of *RAB27A* remained consistent during oocyte maturation, the novel isoform RAB27A-HON-5275 associated with ATTS and elongating 3’ UTR, showed preferential usage at the MII stage (Fig. 4H). Together, these findings implicate isoform switching as a putative mechanism that may contribute to oocyte isoform diversification in human and mouse.

### 5. TE co-option drives novel isoform generation and enhances mRNA stability during oocyte maturation

It is understood that TEs are endowed with the capability to transpose (Senft and Macfarlan, 2021). However, while the majority of mammalian TEs have been found to no longer transpose, TE could regulate the gene expression and drive TE-derived gene products, especially in human and mouse oocytes (Franke et al., 2017; Peaston et al., 2004). We performed full-length isoform sequencing on human and mouse oocytes to comprehensively explore the incorporation of TE sequences within transcripts. According to the transcript loci overlapping with a TE, TE-derived isoforms were categorized into transcription end site (TES), transcription start site (TSS), and exonization types (Fig. 5A). We found that the TE-derived isoforms showed the higher distribution in novel isoforms (27.8% in human, 25.6% in mouse), compared to known isoforms (2% in human, 2.6% in mouse) in oocytes (Fig. 5B and Supplementary Table 3). In contrast to the relatively uniform distribution of TES and TSS in both coding and non-coding types according to the GENCODE reference (Supplementary Fig. 5A), TE-derived isoforms in human and mouse oocytes demonstrated differential preferences for TES and TSS (Fig. 5C and Supplementary Table 4). TES was the most predominant type in human oocytes, with 31.2% coding and 68.8% non-coding transcripts; while TSS was the most predominant type in mouse oocytes, in which coding transcripts occupied two thirds (Fig. 5C and Supplementary Table 4). Further analysis showed the ratios of specific TE families incorporated into transcripts in human and mouse oocytes (Fig. 5D,E). In human oocytes, Alu elements, members of the SINE type, predominated as the primary origins of the TES type (22.2%), and constituted 43% of the coding transcripts in TES type (Fig. 5D and Supplementary Fig. 5B). Whereas ERVL-MaLRs, a subset of the LTR class, significantly contributed to the TSS type (38.4%) and exonization type (32.9%) (Fig. 5D). Notably, ERVL-MaLRs accounted for a considerably large fraction of TSS type (78.2%) in mouse oocytes (Fig. 5E and Supplementary Fig. 5C). Besides, we observed that the significant contributions of ERV1 in human oocytes and ERVK in mouse oocytes to the TSS isoform type, along with previously identified ERVL-MaLRs, accounted for over 66% and 90% respectively (Fig. 5D,E and Supplementary Table 5). In summary, ERVL-MaLR and ERV1, belonging to the LTR class, collectively occupied the majority of the TSS type in human oocytes, while ERVL-MaLR and ERVK accounted for a significant proportion of TSS type in mouse oocytes, either in coding or non-coding transcripts (Supplementary Fig. 5B,C and Supplementary Table 5). We observed that certain TE family members were more likely to be incorporated into oocyte transcripts compared to the transcripts in GENCODE (Supplementary Fig. 5D).

**Figure 5.**
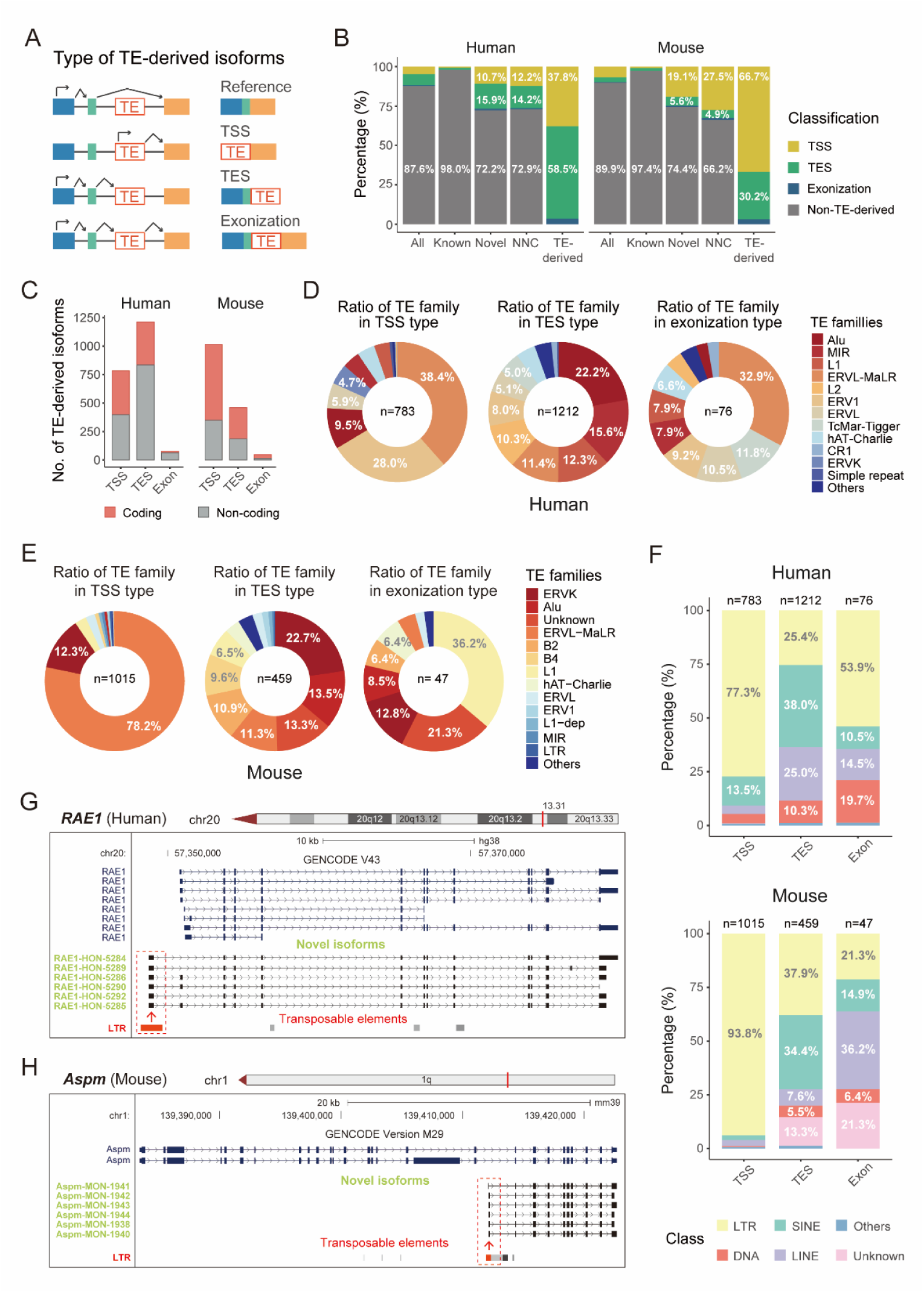
Characteristics of TE-derived isoforms in human and mouse oocytes. (A) Schematic diagram of different types of TE-derived isoforms of the annotation standard. TSS, transcription start site; TES, transcription end site. (B) Percentages of different TE-derived isoform types in all (n = 16,769 in human; n = 14,987 in mouse), known (n = 10,033 in human; n = 10,085 in mouse), novel (n = 6,736 in human; n = 4,902 in mouse), NNC (n = 3,954 in human; n = 2,552 in mouse), and TE-derived isoforms (n = 2,071 in human; n = 1,521 in mouse). (C) The number of different types of TE-derived isoforms in human and mouse oocytes, separated into coding and non-coding transcripts. (D) Ratios of isoforms derived by different TE families in TSS (n = 783), TES (n = 1,212), and exonization (n = 76) types in human oocytes. (E) Ratios of isoforms derived by different TE families in TSS (n = 1,015), TES (n = 459) and exonization (n = 47) types in mouse oocytes. (F) Percentages of TE classes in TSS (n = 783 in human; n = 1,015 in mouse), TES (n = 1,212 in human; n = 459 in mouse), and exonization (n = 76 in human; n = 47 in mouse) types. (G,H) The genomic location and LTR-derived novel isoform structure of *RAE1* in human oocytes (G) and *Aspm* in mouse oocytes (H).

LTR stood out as the most abundant TE class contributing to the TE-derived isoforms belonging to TSS type, comprising 77.3% and 93.8% of the TSS type in human and mouse oocytes, respectively (Fig. 5F). Additionally, LTRs consistently represented the predominant source of the TE-derived isoforms belonging to TSS type in both coding and non-coding transcripts (Supplementary Fig. 5E and Supplementary Table 5). These observations suggested that LTRs were the major contributor to the TSS type, consistent with the known effect of LTRs containing promoters (Hashimoto et al., 2021; Modzelewski et al., 2021). For example, *RAE1* in human oocytes and *Aspm* in mouse oocytes, had multiple novel isoforms belonging to NNC type, and their novel first exons were entirely within the LTR sequence (Fig. 5G,H). RAE1 and ASPM were reported to play key roles in regulating oocyte maturation (Chen et al., 2021; Xu et al., 2012), and the highly expressed novel isoforms derived by LTRs may have important functions during oocyte maturation (Fig. 5G,H and Supplementary Fig. 5F,G). These findings demonstrate the contributions of TEs to the novel isoform generation in human and mouse oocytes.

To explore the widespread influence of TEs across oocyte isoforms, we next analyzed the incorporation of highly expressed TEs within the transcripts in human and mouse oocytes. The most highly expressed TE classes in both human and mouse oocytes were LTR, SINE, LINE, and DNA (Fig. 6A,B and Supplementary Table 3). We have identified that certain family members of these four TE classes could be incorporated into multiple transcripts, rather than providing TE-derived sequences to only one transcript (Fig. 6C). Notably, family members of the LTR class exhibited the highest frequency of incorporation within multiple transcripts in human and mouse oocytes (Supplementary Fig. 5H). Additionally, this capacity for multiple transcripts incorporation was predominantly observed in the TSS type among various TE-derived isoform types (Supplementary Fig. 5I). Our previous findings could exemplify these multi-incorporation events of TEs, where LTRs could supply sequences for multiple novel isoforms of the *RAE1* gene in human oocytes and the *Aspm* gene in mouse oocytes (Fig. 5G,H).

**Figure 6.**
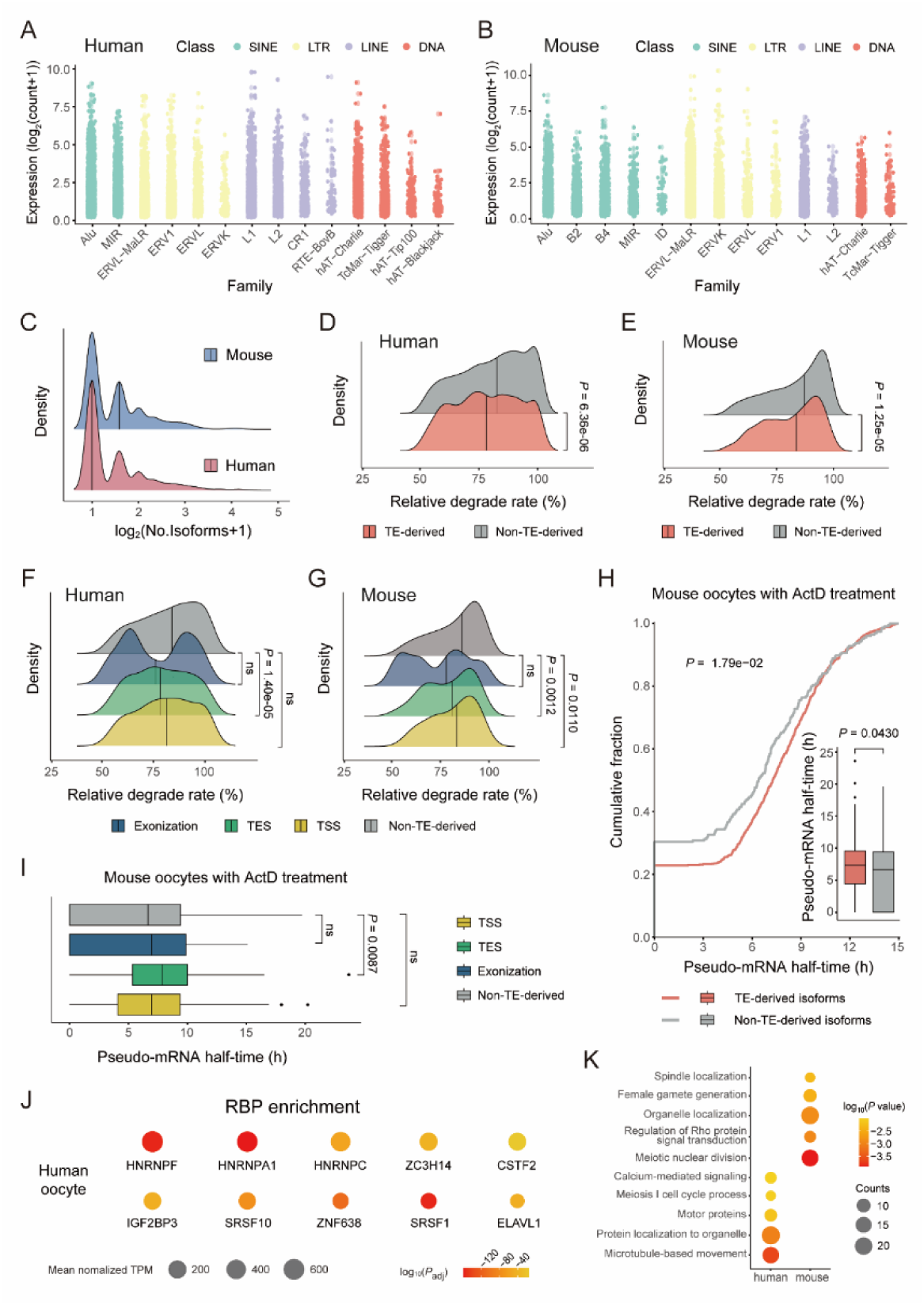
The incorporation of TE sequences within transcripts enhances the mRNA stability during oocyte maturation. (A) Jitterplot of expressed TEs in human oocytes by TE families. The plot was colored by TE classes. TE classes (n > 500) and families (n > 60) are shown. (B) Jitterplot of expressed TEs in mouse oocytes by TE families. The plot was colored by TE classes. TE classes (n > 300) and families (n > 60) were shown. (C) Density plot showing the number of transcripts that individual TE can be incorporated into in human and mouse oocytes. (D,E) Comparison of maternal degradation rates between TE-derived and non-TE-derived isoforms during oocyte maturation in human (D) and mouse (E). (F,G) Comparison of maternal degradation rates between TSS, TES, exonization, and non-TE-derived isoforms in human (F) and mouse (G) oocytes. (H) Cumulative and boxplot illustrating the pseudo-mRNA half-life of TE-derived and non-TE-derived transcripts in mouse oocytes. *P* values were determined by two-tailed unpaired Wilcoxon’s test (boxplot) and K-S test (cumulative plot). (I) Boxplot displaying pseudo-mRNA half-life of TSS, TES, exonization and non-TE-derived isoforms. (J) Dot plots showing the RBP enrichment of TE-derived exons of TES type. Colors indicate significance of enrichment, and dot size represents expression levels of corresponding RBPs. Testing for motif enrichment was conducted using one-tailed Fisher’s exact test with Bonferroni corrected. (K) Enrichment analysis of TE-derived isoforms. The colors show the significance of enrichment; the size of the dots represents the number of genes enriched in each term. Two-tailed unpaired Wilcoxon test was used in D-G, I. Adjustment for multiple comparisons with the Benjamini-Hochberg FDR method was applied in F,G,I. ActD, actinomycin D; RBP, RNA-binding protein.

We observed that the expression levels of TE-derived isoforms were not significantly different from non-TE-derived isoforms in human oocytes (Supplementary Fig. 5J). In mouse oocytes, there was no consistent trend indicating higher expression in either TE-derived or non-TE-derived isoforms (Supplementary Fig. 5K). Moreover, we found that the TE-derived transcripts exhibited a significantly lower maternal degradation rate than that of non-TE-derived isoforms in both human and mouse oocytes (Fig. 6D,E). We also compared the degradation rates between various types of TE-derived isoforms and non-TE-derived isoforms, finding that the TES type displayed a significantly lower degradation rate than non-TE-derived isoforms in both human and mouse oocytes (Fig. 6F,G). The lower degradation rates of TES type predominantly contribute to the differences in stability between TE-derived and non-TE-derived isoforms, suggesting that TEs incorporated into the end site of transcripts might enhance transcript stability during oocyte maturation. To further validate the stability of TE-derived isoforms, we collected mouse oocytes treated with actinomycin D (ActD) and calculated the pseudo-mRNA half-life of transcripts. This assay showed that TE-derived isoforms exhibited a longer half-life than their non-TE-derived counterparts (Fig. 6H). Notably, the TES type displayed a longer pseudo-mRNA half-life, reinforcing the finding on the enhanced stability of TES type compared to non-TE-derived isoforms (Fig. 6I). Further analysis of RNA-binding protein (RBP) enrichment showed that TE-derived exons of the TES type might interact with hnRNP A1 (Fig. 6J), which has been reported to play important roles in mRNA stability (Wang et al., 2016). Enrichment analysis of TE-derived isoform demonstrated that the host genes of these isoforms participate in governing protein localization and meiotic nuclear division (Fig. 6K). These results suggest the potential roles of TEs in bolstering transcript stability and regulating oocyte maturation.

### 6. The NNC isoforms of *ARHGAP18* are specifically expressed and essential during oocyte maturation

ARHGAP18, belonging to the RhoGAP family of proteins, is known to regulate cellular morphology and motility of tumor and endothelial cells (Lovelace et al., 2017; Maeda et al., 2011). In our study, novel isoforms of *ARHGAP18* categorized as NNC type, were identified and found to be specifically and highly expressed in human oocytes (Fig. 2G and Supplementary Fig. 6A). These findings motivated us to further explore the functional significance of *ARHGAP18* during oocyte maturation. The real-time quantitative PCR (RT-qPCR) result showed that novel isoforms of *ARHGAP18* were highly and stably expressed throughout oocyte maturation, while the classical *ARHGAP18* isoforms were almost undetectable (Fig. 7A).

**Figure 7.**
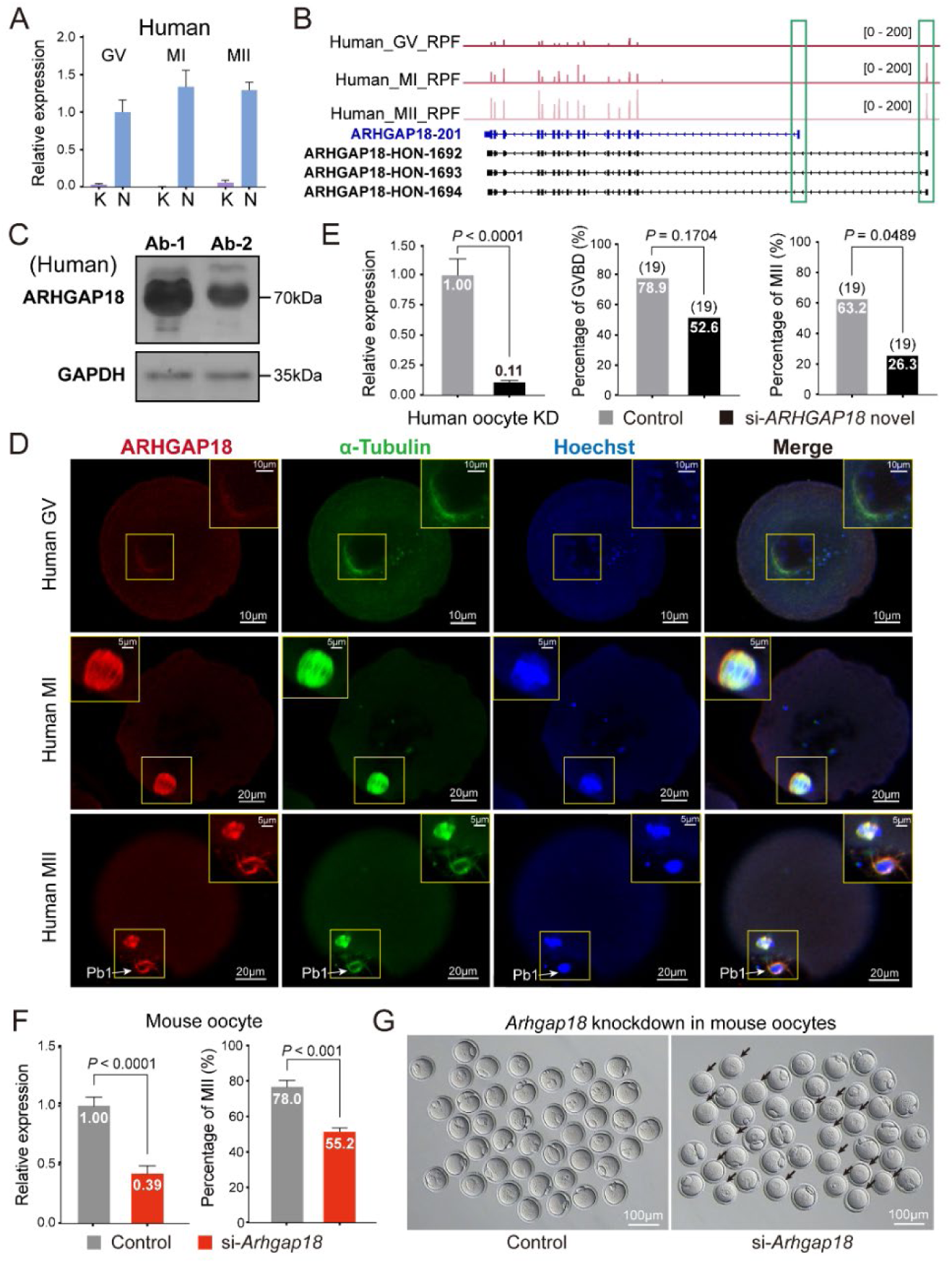
Novel isoforms of *ARHGAP18* are highly expressed and involved in oocyte maturation. (A) Bar graph shows the relative expression of known and novel isoforms of *ARHGAP18* during human GV (n = 14), MI (n = 14), and MII (n = 17) maturation stages through real-time quantitative PCR (RT-qPCR). (B) The ribosome-protected fragment (RPF) signals showing the translation patterns of known and novel isoforms of *ARHGAP18* using published translatomic data. The green box emphasizes the translational expression of novel isoforms of *ARHGAP18*. (C) Western blot demonstrating the expression of the ARHGAP18 protein in human oocytes and early-stage embryos. The protein coded by the novel isoform of *ARHGAP18* is 72.9 kDa as predicted. Ab-1, antibody 1; Ab-2, antibody 2. (D) Immunofluorescent staining showed the co-localization of ARHGAP18 (red) and α-tubulin (green) during oocyte maturation. DNA (blue) was stained with Hoechst. Pb1, first polar body. (E) *ARHGAP18* novel isoform targeted knockdown in human oocytes. The targeted knockdown efficiency was verified by RT-qPCR (left) (n = 15 oocytes for the control group, n = 16 oocytes for the knockdown group), *P*<0.0001, two-tailed Student’s t-test. The percentage of GVBD (middle), *P* = 0.1704, Fisher’s exact test. The percentage of mature MII oocytes (right), *P* = 0.0489, Fisher’s exact test. The number of oocytes analyzed is indicated in parentheses. (F) The knockdown efficiency of *Arhgap18* in mouse oocytes was verified by RT-qPCR (n = 19 oocytes for the control group, n = 25 oocytes for the knockdown group) (left), *P* < 0.0001. The percentage of mature MII oocytes between control and knockdown groups (n = 207 oocytes for the control group, n = 214 oocytes for the *Arhgap18* knockdown group) (right), *P* < 0.001, two-tailed Student’s t-test. (G) Oocyte morphological imaging of the control and knockdown groups (black arrows indicate the immature oocytes within the *Arhgap18* knockdown group). The error bar was shown as mean ± SEM in A,E and F.

At the translational level, a re-analysis of the human oocyte translatome data (Zou et al., 2022) further confirmed the protein coding probability of novel isoforms, and signals for ribosome-protected mRNA fragments increased following human oocyte maturation, suggesting increased protein translation in the MI and MII stages (Fig. 7B). With the exception of the starting 15 amino acid (aa) residues and starting 38 aa residues at the N-terminus of the protein encoded by the respective novel and canonical isoforms, the remaining sequences were identical (Supplementary Fig. 6B). Furthermore, we performed Western blotting and detected an immunoblotted protein band of approximately 70 kDa in human oocytes and cleavage embryos that corresponds with the predicted size of 72.9 kDa (Fig. 7C and Supplementary Fig. 6B). The evidence from both transcriptional and translational levels, and detection of ARHGAP18 protein in oocytes, indicates that the novel isoforms are the major expressed and functional transcripts of *ARHGAP18* in human oocytes.

Immunofluorescence experiments showed that ARHGAP18 localized to the outside of germinal vesicle at GV stage (Fig. 7D). Then, at the resumption of meiosis, ARHGAP18 was co-localized with α-tubulin within the meiotic spindle region from the MI stage to the MII stage, suggesting its role in meiotic spindle assembly in human oocyte. To investigate the roles of *ARHGAP18* novel transcripts in human oocyte maturation, siRNA was designed to specially target the unique exon of novel isoforms for selective knockdown. After microinjection of targeting siRNAs, nearly 90% of *ARHGAP18* novel transcripts were effectively eliminated in human oocytes as confirmed by RT-qPCR, when compared to control (Fig. 7E). Moreover, when compared to control, the siRNA knockdown group showed a decreasing trend of GVBD rate, and a significant reduction in first polar body (Pb1) extrusion (26.3% vs 63.2%, respectively; Fisher’s exact test, *P* = 0.0489) (Fig. 7E). Therefore, these results indicate that these specific novel isoforms of *ARHGAP18* are necessary for oocyte maturation. ARHGAP18 showed high evolutionary sequence conservation among *Homo sapiens*, *Macaca mulatta*, *Sus scrofa domesticus*, and *Mus musculus* (Supplementary Fig. 6C). Further siRNA-mediated knockdown of *Arhgap18* in mouse oocytes showed a significant decrease in Pb1 extrusion (*P* < 0.001, two-tailed Student’s t-test) (Supplementary Fig. 6D, Fig. 7F, G). Taken together, our results in human and mouse oocytes demonstrate that ARHGAP18 plays a conserved and vital role in oocyte maturation.

## Discussion

Here, we have developed single oocyte long-read sequencing to characterize full-length transcripts and generate detailed isoform level atlases of transcriptome over the course of human and mouse oocyte maturation. Considerable isoform diversity was identified among oocyte maturation stages across both species, with about 40% of which characterized as novel transcripts not annotated according to the GENCODE reference, which is distinctly higher than that in GCs and ESCs. By generating reads spanning entire transcripts, a total of 6,736 and 4,902 novel isoforms were characterized in human and mouse oocytes, respectively. Comparing our data with assemblies of previously published studies and GENCODE database, we found that around 36% of the full-length isoforms identified were previously undetected (Franke et al., 2017; Hu et al., 2022; Liu et al., 2019; Llonch et al., 2021; Zhang et al., 2018; Zhao et al., 2020). Our full-length isoform sequencing provides a relatively comprehensive isoform-level transcriptome profile, underscoring its capacity to detect novel isoforms. Notably, the vast majority of novel transcripts were characterized as NNC in oocytes of both species, markedly different to that in other cell types like the cerebral cortex, cancer, and cell lines, where NIC transcripts comprised the dominant component of novel transcripts (Leung et al., 2021; Liao et al., 2023; Veiga et al., 2022). It suggests that the isoform spectrum similarity of the homologous cell types, such as the oocyte, in different species is greater than that of different cell types in the same species.

We discovered NNC category isoforms of *ARHGAP18* which were abundant and specifically expressed in human oocytes. The known isoforms of *ARHGAP18* are almost not expressed in oocytes, hence its function in oocytes has not been explored. The existence and microtubular localization of ARHGAP18 in human oocytes, and the targeted knockdown of its novel isoforms results in oocyte maturation defects, implying the crucial functions of *ARHGAP18* in meiotic spindle assembly and stability. We believe that the identification of oocyte-specific transcripts with unannotated regions or splicing variants through full-length isoform sequencing, would expand the potential risk genes relevant to clinical diagnosis of oocyte maturation defect.

In addition to identifying novel transcripts, we also characterized the DTU events during oocyte maturation in human and mouse oocytes. Although there is global transcriptional silencing accompanied by degradation in total mRNA levels during oocyte maturation, some DTU events showed increased isoform fraction at MII stage during oocyte maturation at isoform level. In both species, a large proportion of transcripts underwent ATTS gain events and showed a trend of 3’ UTR elongation during maturation. These findings suggest that AS could serve as a putative mechanism for DTU events. Additionally, partial DTU events may result from differential degradation rates among the existing isoforms.

More than a quarter of the novel isoforms we identified have been annotated as TE-derived isoforms, which suggests that TEs provide important origins to generate novel isoforms during mammalian oocyte maturation. Previous research has demonstrated that ERVL-MaLRs have been co-opted as alternative promoters for adjacent genes (Flemr et al., 2013). In addition to ERVL-MaLRs, we found that ERV1 elements also constitute a significant proportion of TSS type in human oocytes, suggesting the potential roles of ERV1 in supplying promoters for oocyte-specific isoforms crucial for oocyte maturation. Additionally, the greater stability of TES type primarily contributed to the differences in stability between TE-derived and non-TE-derived isoforms, implying that the incorporation of TE sequences within the end site of transcripts might bolster isoform stability during oocyte maturation.

In summary, we comprehensively profiled the isoform-level mapping of oocyte maturation and identified high-confidence potential novel genes and isoforms, providing a valuable resource for designating the molecular framework of oocyte maturation. Our results highlighted that TE-derived isoforms are an important source of oocyte-specific isoforms, and pre-existing TE sequences could be co-opted into oocyte transcripts, potentially contributing to mRNA stability during oocyte maturation. Thus, our findings add and expand upon the gene and transcript annotation available for the developmental biology community and point to a potential mechanism for oocyte maturation involving the enhanced stability of TE-derived isoforms.

## Methods

### 1. Ethics statement

The research proposal for experiments with clinically discarded human oocytes and GCs was reviewed and approved by the Reproductive Medicine Ethics Committee of Peking University Third Hospital (Approval No. 2019SZ-028, M2023617). The committee was composed of 14 members with expertise in reproductive medicine, psychology, ethics, jurisprudence, sociology, clinical epidemiology and general public. The research approach, including criteria for oocyte collection and manipulation, and the informed consent process were all assessed by the committee. Informed consent was obtained from the participants before sample collection. The investigators of this study took charge to explain the research objectives, the implications of donations and potential risks involved, to ensure clarity and to support informed decision-making. Each participant provided written informed consent for the collection of oocytes for research purposes voluntarily, without receiving any compensation. Human oocytes and GCs were acquired from women (aged 22 to 34) who went to Peking University Third Hospital due to male factor infertility (human oocytes obtained from September 2021 to June 2022, and human GCs obtained from June to July 2024). All human oocytes employed in this study were not viable for clinical use and have not been genetically modified.

Experiments with mouse oocytes and GCs were conducted according to protocols approved by the Ethical Committee of Laboratory Animals (Approval No. LA2019341, A2024075) and was compliant with all relevant ethical regulations regarding animal research. All mice were bred and maintained according to institutional guidelines. All experiments followed the 2021 Guidelines for Stem Cell Research and Clinical Translation released by the International Society for Stem Cell Research (ISSCR).

### 2. Collection of human and mouse oocytes

For experiments with human samples, only the oocytes that were in an immature state during in vitro fertilization (IVF) or intracytoplasmic sperm injection (ICSI) were collected and used. Human oocytes used for sequencing were acquired from 20 female patients ranging from 22 to 34 years old who underwent IVF or ICSI due to male factor infertility. As a quality control measure for human oocytes used in this study, only oocytes with normal morphology that underwent GVBD and maturation within 24-48 hours after oocyte retrieval operation and in vitro maturation (IVM) procedures (Li et al., 2008; Liu et al., 2021) were collected and used. Samples from females with medical conditions such as obesity and polycystic ovary syndrome were excluded.

For mouse experiments, mice were housed in a specific pathogen-free facility with a 12 hours light/dark cycle, 50 % relative humidity, and an ambient temperature of 22 ℃. To collect GV oocytes, 8- to 10-week-old B6D2F1 female mice were injected with 10 international units of pregnant mare serum gonadotropin (NSHF, China). Forty-eight hours later, oocytes were isolated by ovarian puncture and collected by mouth pipette. The harvested GV oocytes were divided into a pool for direct collection and processing, as well as a pool that was cultured in M2 medium (Sigma-Aldrich, USA) for IVM. MI oocytes were collected 6 hours after GVBD, and MII oocytes were collected after extrusion of Pb1. Before their collection, all oocytes were exposed to a solution of 3‰ hydrochloric acid for ten seconds to remove the zona pellucida followed by three washes in PBS.

### 3. Oocyte lysis and cDNA preparation

Each individual oocyte was manually picked and transferred into a tube containing 4 μL of lysis buffer with 1×10^4^ External RNA Controls Consortium (ERCC) (Thermo Fisher, USA) and stored at −80 °C until use. Samples were vibrated sufficiently and incubated in 72 °C for 3 min to release the RNA into the lysate (Picelli et al., 2014). Next, the lysate was subjected to reverse transcription and pre-amplification using a SMART-seq2 kit according to the published protocol (Picelli et al., 2014). During the process of reverse transcription, barcoded primers containing poly T, UMIs, barcodes as well as PCR oligo sequences, were ligated to the cDNAs. Distinct UMIs were ligated to each cDNA molecule to eliminate the bias from PCR amplification, while unique barcodes were ligated to cDNAs from individual oocytes to facilitate the subsequent pooling process. Prior to library synthesis, both the quantity and quality of cDNA were assessed using a Qubit™ fluorometer (Thermo Fisher, USA) and Fragment Analyzer (Agilent Technologies, USA), respectively.

### 4. Full length isoform sequencing (Iso-Seq) library preparation and SMRT PacBio sequencing

A total of 20 human oocytes and 26 mouse oocytes were pooled to attain the necessary 500 ng input quantity for library preparation. The pooled cDNA was synthesized into SMRTbell libraries using a SMRTbell Express Template Prep Kit 2.0 (Pacific Biosciences, USA), according to the manufacturer’s protocol. After purification, SMRTbell cDNA libraries were loaded and sequenced using the CCS mode of a PacBio Sequel II System to generate HiFi reads.

### 5. RNA-seq library preparation and Illumina sequencing

Approximately 250 ng of each amplified cDNA was used for the subsequent fragmentation procedures. Samples were fragmented to produce cDNAs of approximately 300 bp in length using a Covaris Ultrasonicator. The fragmented cDNA was then used to generate sequencing library using a NEBNext® Ultra™ II DNA Library Prep Kit for Illumina® (New England Biolabs, USA) according to the manufacturer’s protocol. Sequencing libraries were purified by AMPure XP beads (Beckman, USA) and DNA concentrations were assessed using Qubit™ fluorometer (Thermo Fisher, USA). An Illumina NovaSeq 6000 platform was used for library sequencing.

### 6. Collection and sequencing of human and mouse ESCs or GCs

Primed H9 human ESCs were kindly provided by Professor Jie Na (Tsinghua University). Primed H9 ESCs were cultured on Matrigel® Matrix-coated plates (Corning, USA) with complete TeSR-E8 Basal medium (Stem Cell Technologies, Canada). During cell passaging, they were dissociated using Accutase (Stem Cell Technologies, Canada) and resuspended with medium containing ROCK inhibitor (Y-27632) (Selleck Chemicals, USA). ES-E14TG2a (E14) mouse ESCs were purchased from ICell Bioscience Inc (Shanghai, China). Cells were cultured with mESC medium consists of 81% DMEM high glucose (Gibco™, USA), 15% fetal bovine serum (Gibco™, USA), 1% GlutaMAX-1 (Invitrogen, USA), 1% MEM NEAA (Invitrogen, USA), 1% Sodium Pyruvate (Gibco™, USA), 1% Penicillin-Streptomycin (Gibco™, USA), 1× LIF (Millipore, USA) and 1× β-Mer (Invitrogen, USA). Both human H9 and mouse E14 ESCs were incubated at 37℃ under 5% CO_2_.

Human GCs were acquired from female patients who received ICSI due to male sterile factors. Human GCs were mechanically separated from cumulus oophorus complex (COC) and washed in PBS for at least three times before collected into RNA lysis. Mouse GCs were acquired from 8-10-week B6D2F1 female mice. COCs were obtained using the method described above. Then, the GCs were digested to single cells using hyaluronidase and washed in PBS for at least three times. Human and mouse GCs were separately collected into RNA lysis, with 10 cells per tube.

A total of 16 samples (4 human H9 ESCs, 4 human GCs, 4 mouse E14 ESCs, and 4 mouse GCs) cDNAs were pooled together to attain the necessary 500 ng input quantity for Iso-Seq library preparation. The Iso-Seq library preparation and SMRT PacBio sequencing procedures were same as oocyte samples mentioned above. The RNA-seq library construction and Illumina sequencing procedures used for human and mouse ESC and GC samples were same as oocytes.

### 7. Full-length isoform sequencing data processing

Raw subreads were processed with the Iso-seq3 (3.4.0) pipeline (https://github.com/PacificBiosciences/IsoSeq). In brief, ccs module was used to call CCS reads, including reads with an accuracy over 90% (‘--min-rq 0.9’). Lima (2.4.0) was used to orient demultiplex reads based on cell barcodes followed by UMI tagging (--design T-8U) and poly(A) tail trimming of tagged full-length reads. Duplicates were removed to produce deduplicated bam files.

### 8. TGS Transcriptome annotation

The refined, full-length tagged non-concatemer (FLTNC) reads were clustered using the ‘isoseq3 cluster’ function and aligned to reference genomes (GENCODE hg38 v40 for human; mm39 vM29 for mouse) using pbmm2 (v1.9.0). Transcript structures were collapsed using cDNA_cupcake to get the representing isoform with the longest 5’ ends.

Classification was performed using SQANTI3 (v4.2), which categorized isoforms as FSM, ISM, NIC, or NNC based on splice sites and exon structure. An isoform is classified as FSM if it fully aligns with the reference genome, has the same splice junctions, and contains the same number of exons. If it contains fewer 5 prime exons compared to the reference isoform, it is classified as an ISM. A novel isoform that contains a combination of known donor or acceptor sites is classified as a NIC, while a novel isoform with at least one novel donor or acceptor site is classified as an NNC. The classification criteria and schematic description of each isoform category are shown in Figure 2A.

To predict the coding probability and the open reading frame length of the isoforms, CPAT (3.0.4) (Wang et al., 2013) was utilized with default parameters. Isoforms were classified as protein-coding if the coding probability score was greater than or equal to 0.364 for humans and greater than 0.44 for mice, according to a default parameter (Wang et al., 2013).

We included the isoforms that exist simultaneously in at least three samples for subsequent analyses. Novel isoforms that were not FSM or ISM were defined as human oocyte novel (HON) or mouse oocyte novel (MON) isoforms. Gffread (0.12.7) (Pertea and Pertea, 2020) was used to convert gtf files into fasta files.

The same pipeline used for oocyte samples was applied to Iso-Seq data from ESC and GC samples for calling CCS reads, demultiplexing, refining and deduplicating. For the transcriptome assembling of ESC and GC, only isoforms present in at least two biological replicates were retained.

### 9. Quality control and preprocessing of RNA-seq data based on NGS

Raw fastq data from NGS platform was trimmed by Trim Galore (0.6.6) with the parameter of ‘--paired --quality 20 --phred33 --stringency 3 --length 36’. The trimmed fastq files were then mapped to reference genomes (mentioned above) using STAR (2.7.8a), with the parameter ‘--readFilesCommand zcat --outSAMtype BAM SortedByCoordinate --outReadsUnmapped Fastx’. The isoform expression matrices were obtained by salmon (1.9.0) (Patro et al., 2017). Stringtie (2.1.1) was used to assemble transcriptome gtf files. Quality control was performed based on the number of detected genes, uniquely mapping rates, and the percentage of reads mapping to mitochondrial genes, and only samples that passed the quality control were retained according to a previously published study (Zhang et al., 2018). Total 20 human oocytes and 26 mouse oocytes were retained for the downstream analysis.

### 10. Comparison of RNA-Seq and Iso-Seq data

SQANTI3 was used to compare assemblies of RNA-Seq and Iso-Seq data (parameters: --CAGE_peak --polyA_motif_list). The comparison provides quality features including annotation on independently published Cap Analysis of Gene Expression (CAGE) peak and 3’ ends of poly A motifs (Forrest et al., 2014; Pardo-Palacios et al., 2024).

### 11. Quantification of isoform expression

Deduplicated full-length reads were mapped to the reference transcriptome we assembled using minimap2 (2.17) (Li, 2018) (parameters: -t 30 -ax splice -uf --secondary=no -O6,24 -B4). Mapped reads were quantified with salmon (1.9.0) (parameters: -l U --minAssignedFrags 1 –noErrorModel) to obtain isoform expression matrices. NanoStat (1.6.0) was used to create statistical summaries of all long reads. RSeQC (2.6.4) (Wang et al., 2012) was used to evaluate gene body coverage.

### 12. Differential analysis at gene and isoform level

All differential analyses are based on TGS data. For DEGs, the gene expression matrix generated by aggregating the counts of all isoforms of each gene, was analyzed in DESeq2 (Love et al., 2014) with ERCC spike-ins as size factor for normalization. Genes with average transcripts per million (TPM) > 5, adjusted *P* value (FDR) < 0.05, and log_2_ fold change > 1 were considered differentially expressed.

For differential transcript expression (DTE), the isoform expression matrix was filtered to include only isoforms with isoform fraction (IF, calculated as the isoform expression divided by the gene expression) > 0.01, and host gene average TPM > 1. After filtering, the same parameters and cutoff criteria for DEGs analysis were applied. Only transcripts from genes with multiple isoforms were finally considered as DTE (Patowary et al., 2024).

DTU refers to the changes in the proportions of the various isoforms of a gene, and was performed using DEXSeq (Li et al., 2015) as implemented in IsoformSwitchAnalyzeR (Vitting-Seerup and Sandelin, 2019). The normalized isoform counts matrix was extracted from DESeq2 following the same parameters used in DEGs and DTE analyses. For DTU, the difference in isoform fraction (dIF) value between two stages and FDR adjusted *P* value were calculated. Isoforms with dIF > 0.1 and adjusted *P* value < 0.05 were considered to exhibit significant DTU. At gene level, any gene with at least one significant isoform was considered to exhibit differential transcript usage.

### 13. Enrichment analysis

Enrichment analyses were conducted using Metascape (Zhou et al., 2019) with the default parameters. Ggplot2 was used to visualize the results. The biology process related genes were obtained from Metascape.

### 14. Analysis of ribosome-protected mRNA fragments (RPFs)

Raw sequencing data for RPFs was obtained from a previously published study (Zou et al., 2022). Cutadapt (3.3) was utilized to trim adaptors and primers from the sequenced reads. Bowtie2 (2.4.2) (Langmead and Salzberg, 2012) was used to remove the reads mapped to ribosome RNA with the parameter ‘--seedlen=23’. Filtered bam files were then mapped to the human genome using STAR (2.7.8a) (Dobin et al., 2013). Bedtools multicov (2.30.0) (Quinlan and Hall, 2010) was utilized to count the coverage of RPFs at specific genomic loci. BamCoverage (3.5.1) (Ramirez et al., 2014) was used to convert the bam files into bigwig files for visualization of coverage through the Integrative Genomics Viewer (IGV) (Robinson et al., 2011).

### 15. Transposable element expression analysis

RepeatMasker sequences and annotations (human: hg38; mouse: mm39) were downloaded from the UCSC table browser for TE associated analysis. Deduplicated full-length reads were mapped and quantified to RepeatMasker sequences and annotations with minimap2 (2.17) and salmon (1.9.0).

### 16. Annotation and expression analysis of TE-derived isoforms

The assembled transcripts mentioned above were utilized in the annotation of TE-derived isoforms in human (n = 16,769) and mouse oocytes (n = 14,987), respectively. The annotation pipeline of TE-derived isoform was adapted and modified as previously described (Berrens et al., 2022), summarized briefly as follows:

1. If an exon of the queried isoform does not overlap 100% with any known exon loci from GENCODE database and it has over 100nt overlap with the locus of any TE, this exon is annotated as a TE-derived exon.
2. If the 5’ exons of the queried isoform are TE-derived exons, the annotation designates the isoform as TSS. Conversely, when the 3’ exons of the queried isoform are TE-derived exons, the isoform is categorized as TES. Where any internal exon of the queried isoform is TE-derived, while all other exons remain non-TE-derived, the isoform is annotated as exonization.

The annotation of TE-derived isoforms in GENCODE V40 and GENCODE VM29 assemblies followed the same methodology applied to our assembled transcripts. Following the annotation of all TE-derived isoforms, deduplicated bam files were mapped to our transcript assemblies with minimap2 (2.17). Salmon (1.9.0) was then used to quantify the expression of isoforms.

### 17. Analysis of maternal degradation transcripts

As described in previous research (Sha et al., 2020), maternal transcripts that were detected in at least three samples and with a TPM of >2 at the GV stage were retained for the maternal degradation analysis. Isoforms were defined as maternal degradation isoforms that satisfied the following criteria: log_2_ (TPM of GV+1) > log_2_ (TPM of MII+1) + 1. The relative degradation rate was then calculated as (TPM of GV-TPM of MII) /TPM of GV×100%.

### 18. Analysis of publicly available deepCAGE and PAIso-seq data

The deepCAGE data analysis was performed as previously described (Babarinde et al., 2021). Published deepCAGE bam files of human and mouse ovaries were collected from FANTOM5 databases (Pertea and Pertea, 2020). PAIso-seq data during human and mouse oocyte maturation were obtained, and the preprocess was performed as previously described (Liu et al., 2019; Liu et al., 2023).. BamCoverage (3.5.1) was utilized to generate the bigwig files. ComputeMatrix (3.5.5) module of deeptools was used to calculate reads density (parameters: scale-regions -b 2000 -a 2000). CrossMap (0.7.0) was used to liftover the bigwig files from mm9 into mm39.

### 19. Analysis of publicly available RNA-seq data

Published RNA-seq data during human and mouse oocyte maturation were collected (Franke et al., 2017; Hu et al., 2022; Liu et al., 2019; Llonch et al., 2021; Zhang et al., 2018; Zhao et al., 2020). Adapters, PCR oligo and poly A tails were trimmed by house-in scripts or Trim_Galore (0.6.6) according to the structure of libraries. STAR (2.7.8a) was used to align the clean reads to GENCODE reference. Stringtie (2.1.1) was used to assemble transcriptome gtf files of merged bam files. Gffcompare (v0.11.2) was utilized to assess the transcript similarity across different datasets, considering two transcripts as identical when they exhibited an exact intron chain match, according to a previously published study (Patowary et al., 2024).

### 20. The validation of novel transcripts by Sanger sequencing following PCR

The authenticity of novel transcripts was validated by analyzing the rest of cDNAs from sequenced oocyte and cDNAs from other oocytes. Isoform-specific primers were designed using the Genome Browser website (UCSC: https://genome.ucsc.edu) and PrimerQuest™ Tool (https://sg.idtdna.com), covering the unique exons of the novel transcripts. After amplification of the target isoforms using OneTaq® Hot Start 2X Master Mix (New England Biolabs, USA), the amplicons were separated by electrophoresis on a 1% agarose gel and detected by Sanger sequencing. To accurately verify the authenticity of ISM transcripts, the amplified fragments of ISM transcripts were inserted into a T vector before Sanger sequencing. The primer sequences used were listed as below: *ELOVL5*-ISM-2 5’ region, forward primer: ATGTACCCGTTTGTGATGAACT, reverse primer: CAGAGAGCCCAGAAATGTTAGA; *EBLN1* novel isoforms, forward primer: CTCTTTCTCTAGGTCCTGCTG, reverse primer: TTTGCTCACACCGACACT; *ARHGAP18* novel isoforms, forward primer: GGCAGCTGCTGGATGAATAA, reverse primer: GGTCCGCGTCAATGTTGATA.

### 21. mRNA Stability Assay

To determine the stability of mRNAs in oocytes, ActD was used to completely inhibit the transcription of mouse oocytes (HY-17559, MCE). GV stage oocytes were obtained from 8–10-week B6D2F1 mice using the aforementioned superovulation and oocyte collecting methods. M2 medium containing 4 μM ActD were used for in vitro oocyte maturation culture. According to previous studies, oocyte samples were collected into RNA lysis (one oocyte per tube) at 0, 2, 4, 6, 8, 10 and 12 hours with 4 replicates at each time point and group. The reverse transcription and amplification, RNA-seq library preparation and sequencing procedures were same as described in this study.

The pseudo-mRNA half-life analysis was performed as previously described (Li et al., 2024). RNA-seq data were processed using fastp (0.23.4) for adaptor trimming and quantified with Salmon (1.9.0). The RNA degradation rate k_decay_ was defined as the mean values of ratio from reads per kilobase per million mapped reads (RPKM) at time 0 hour over the RPKM value at each other time point (2, 4, 6, 8, 10 and 12 hours). The pseudo-mRNA half-life of the genes expressing both TE-derived and non-TE-derived transcripts t_1/2_ was calculated as ln_2_/k_decay_ as previously described (Li et al., 2024).

### 22. RBP motif enrichment analysis

The TE-derived exons of TE-derived isoforms in the TES type were extracted to perform the RBP motif enrichment analyses. A unified compendium of approximately 1,600 RNA motifs associated with 186 RBPs was compiled according to a previously published study (Hu et al., 2022). AME module of MEME suite (5.4.1) was utilized to perform the RBP motif enrichment with TE-derived exons as target fasta and our assemblies as background fasta with default parameters.

### 23. Real-time quantitative PCR (RT-qPCR)

Each single human oocyte was collected and reverse transcribed into cDNA using the same method as described before in this study. RT-qPCR was performed using PowerUp SYBR Green Master Mix (Thermo Fisher, USA) on a QuantStudio 3 Real-Time PCR System (Thermo Fisher, USA). Relative mRNA levels were calculated by normalizing to the internal control of *ACTB* using the 2^-ΔΔCt^. The primer sequences used were listed as below: *ARHGAP18* known isoform, forward primer: TCCAGTTCCCAGGGAGT, reverse primer: GCCTTTGCATGGCTGTTC; *ARHGAP18* novel isoform, forward primer: GCTGGATGAATAAGCCGAAATG, reverse primer: CTTCCGTGACACACCTACTTT; *ACTB*, forward primer: AGAGCTACGAGCTGCCTGAC, reverse primer: AGCACTGTGTTGGCGTACAG.

### 24. Western blotting

Approximately 100 human oocytes or embryos collected prior to the 8-cell stage were lysed in Laemmli buffer (Bio-Rad, USA) containing 2-hydroxy-1-ethanethiol (Sigma-Aldrich, USA) and heated at 100 ℃ for 5 min. Protein samples were subjected to 8 % SDS-PAGE for separation and transferred to 0.45 μm polyvinylidene fluoride (PVDF) membrane, then blocked in 1×TBST with 5 % nonfat milk for 1 hour. Then, the PVDF membrane was incubated overnight at 4 ℃ with anti-ARHGAP18 polyclonal antibody (1:300 dilution, YT0317, Immunoway, USA; 1:500 dilution, PA5-98757, Thermo Fisher, USA). After several washes in 1×TBST, HRP-conjugated goat anti-rabbit-IgG H&L (1:2000 dilution, ab6721, Abcam, USA) was used as secondary antibody and incubated with the membrane for 1 hour at room temperature. Finally, signals were detected using Pierce™ ECL Western blotting substrate (Thermo Fisher, USA).

### 25. Immunofluorescence

The zona pellucida of human oocytes were removed using 3 ‰ hydrochloric acid solution, followed by three washes in PBS droplets. Next, oocytes were fixed in 4 % paraformaldehyde solution for 30 min at room temperature and then washed with buffer (0.01 % Triton X-100 and 0.1 % Tween-20 in PBS). Oocytes were permeabilized in a solution of 0.3 % Triton X-100 in PBS for 30 min and subsequently blocked in PBS solution with 5 % normal goat serum and 3 % BSA. Next, oocytes were incubated overnight at 4 ℃ with Arhgap18 primary antibodies (1:100 dilution, YT0317, Immunoway, USA) diluted in blocking solution. After washing three times in buffer, oocytes were incubated in secondary Alexa Fluor™ 594 goat anti-rabbit IgG (A32740, Invitrogen, USA), and FITC anti-alpha-tubulin antibody (ab64503, Abcam, USA) for 1 hour at room temperature. After a subsequent wash, oocytes were counterstained using Hoechst 33342 (62249, Thermo Fisher, USA). Oocytes were placed in a confocal dish with anti-bleaching solution (Beyotime, China) and analyzed by confocal microscopy (Zeiss LSM 880).

### 26. Human *ARHGAP18* novel isoforms siRNA knockdown

Human GV stage oocytes with normal morphology were injected with 20 μM *ARHGAP18* siRNA (sequence GCAGCTGCTGGATGAATAA) specifically targeting the unique region of novel isoforms or 20 μM negative control sequence, which were synthesized and provided by the same company (Guangzhou RiboBio Co., China). Injected GV stage oocytes were arrested in M2 medium containing milrinone (Selleck, USA) for 6 hours to allow for mRNA translation, and subsequently released by repeatedly washing oocytes in M2 medium. Then the released human oocytes were incubated in Geri™ Time Lapse under a humidified atmosphere of 5 % CO_2_ at 37 °C for around 32 hours to record the IVM process. The targeted siRNA knockdown efficiency was validated by RT-qPCR.

### 27. Mouse *Arhgap18* siRNA knockdown

Eight-week ICR female mice were used for experiments to knock down *Arhgap18* using targeting siRNAs. Housing of mice, protocols for ovarian stimulation, and GV oocyte retrieval method were all as described above. GV oocytes were injected with 20 μM *Arhgap18* siRNA 1 and 2 (siRNA1, CGCAGATATGGCCAATACA; siRNA2, CAGAATCTTCCAACCAGAA), or negative control sequence. The *Arhgap18* siNRAs and negative control siRNAs were synthesized and provided by the same company (Guangzhou RiboBio Co., China). Injected oocytes were arrested at the GV stage in M2 medium containing milrinone (Selleck, USA) for 16 hours to allow for mRNA translation. Then, the oocytes were washed in at least three clean drops of M2 medium before they were cultured in M2 medium under a humidified atmosphere of 5 % CO_2_ at 37 °C for 20 hours, and the IVM process was recorded by Geri™ Time Lapse. siRNA knockdown efficiency was validated by RT-qPCR. The primer sequences used for mouse *Arhgap18* were listed below, forward primer: GAGCAAGCTGTCAATCAGAAA, reverse primer: CTCCAATCCTTGTCGTACCT.

### 28. Statistics and reproducibility

GraphPad Prism 10 and R (v 4.3.1) software was used to perform all statistical analyses and data visualizations. Experiments were performed with at least three biological replicates. Fisher’s exact test was used to assess the significance of GVBD and MII rates following knockdown of *ARHGAP18* novel isoforms in human oocytes, while other outcomes between the two experimental groups were analyzed with a two-tailed Student’s t-test. For bioinformatics analyses, two-sided rank-sum Wilcoxon tests (non-parametric) were applied, with multiple comparisons adjusted using the Benjamini-Hochberg FDR method. All statistical methods are described in detail in the Methods section, and additional information is provided in each figure legend. *P* value < 0.05 was considered statistically significant throughout the analyses.

## Acknowledgments

We thank the generous donors whose contributions have enabled this research. We thank all the staff in the Center for Reproductive Medicine of Peking University Third Hospital. This project is funded by National Natural Science Foundation of China (82288102, 82125013, 82201838), National Key Research and Development Program (2022YFC2702200, 2023YFA1800300, 2019YFA0801400), Peking University Third Hospital Fund for Interdisciplinary Research (BYSYJC2023001), Clinical Medicine Plus X - Young Scholars Project, Peking University, the Fundamental Research Funds for the Central Universities (PKU2024LCXQ005), and CAMS Innovation Fund for Medical Sciences (2019-I2M-5-001).

## Author contributions

J.Q., P.Y., L.Y., and Q.L. supervised the project. Y.W., W.W., Y.L., Q.L., P.Y., L.Y., and J.Q. conceived and formulated the concept. Y.W. and Y.L. collected the samples and performed all experiments including library construction, single-cell sequencing, siRNA knock down, mRNA stability assay, RT-qPCR, Western blotting, and immunofluorescence under the supervision of P.Y., L.Y., and J.Q. W.W. and Y.H. conducted bioinformatics analyses and visualization on full-length isoform sequencing and next generation sequencing data under the supervision of P.Y. M.Y., N.W., X.W., L.D., Y.K. and Y.L. assisted with samples collection and siRNA injection. Y.X., Z.D., and L.C. assisted with ESCs and GCs collection. Y.W., W.W., Y.L., Y.H., and H.S. drafted the initial manuscript. All authors discussed the results and reviewed the manuscript.

## Data availability

All of the raw sequencing data generated in this study have been deposited in the Genome Sequence Archive for human (GSA-Human) and Genome Sequence Archive of China National Center for Bioinformation (Human: HRA006583; Mouse: CRA014675). The processed data including assembled transcriptomes (annotated gtf files), new splice junctions and expression tables are publicly available in the OMIX, China National Center for Bioinformation (OMIX008807, OMIX008808). Publicly available RNA-seq datasets used in this work were retrieved from SRA (SRP011546, SRP285893, SRP361878, SRP086707, SRP189718, SRP062106). Publicly available deepCAGE and PAIso-seq datasets used in this work were retrieved from FANTOM5, GSA-Human (HRA001288) and SRA under the accession number PRJNA529588.

## Code availability

No original code or algorithm was generated of this study. All analysis utilized publicly accessible algorithms and software. The codes supporting this study can be obtained from the corresponding author upon reasonable request.

## Supplementary information

Supplementary Figures: Supplementary Figures 1-6. Supplementary Tables: Supplementary Tables 1-5.

