## Supplemental_Figures_1_6 for "Single oocyte full-length isoform sequencing unveils the impact of transposable elements on RNA diversity and stability during oocyte maturation"

### Supplementary Figures

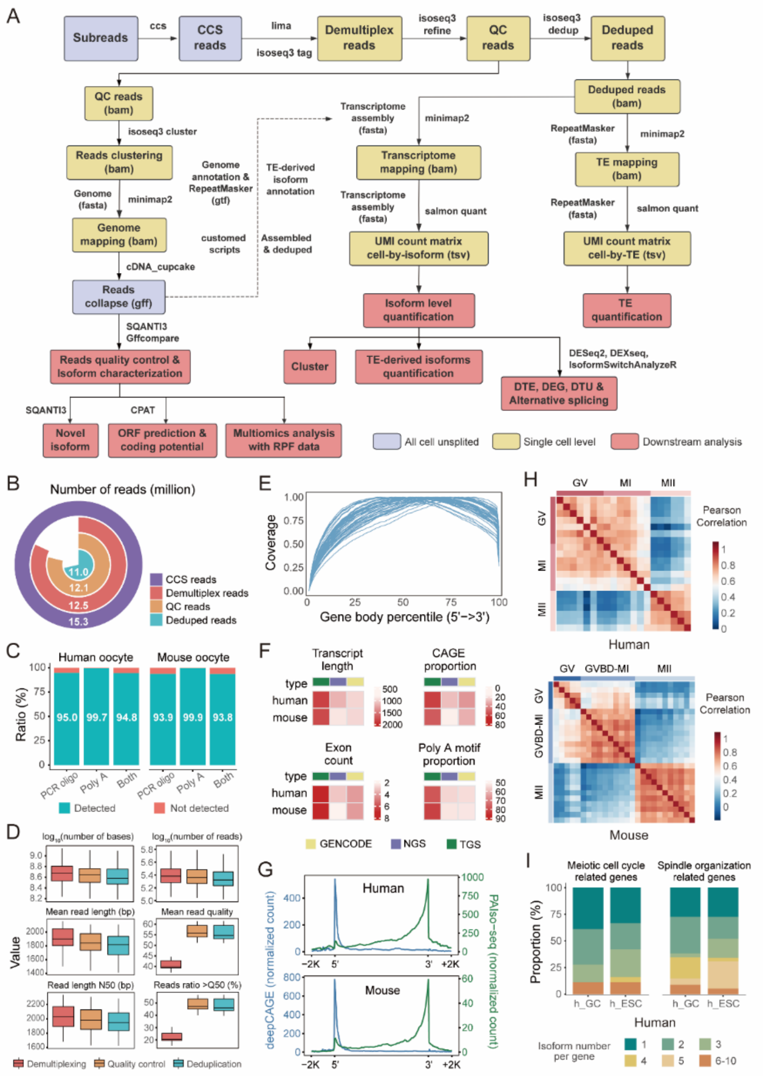


**Supplementary Figure 1. Detailed workflow and quality control protocol for full-length isoform sequencing data.** (A) Flow chart describing the workflow for resolving full-length isoform sequencing data. Steps for calling circular consensus sequence (CCS) reads, demultiplexing, adding tags, trimming, quality control, and deduplication of SMRT subreads are shown above. The nodes are distinguished by their data scale, with yellow representing single-cell level, red representing tab-separated matrix, and purple representing all cells unsplit. The file format for each node is indicated within brackets. (B) Rose plot depicts the number of reads after each processing step. (C) Ratio of demultiplex reads with PCR oligo and poly A tails. (D) Boxplots display statistical metrics for yield, length, and quality of all TGS data after demultiplexing, quality control, and deduplication procedures for each individual cell. (E) Genebody coverage of reads along the whole gene body of full-length isoforms. (F) Pileup heatmaps comparing the quality of transcripts between transcriptome assembled using TGS and NGS, along with GENCODE transcripts, including median transcript length (excluding introns), median exon number, CAGE and poly A support for isoforms. (G) Read density of ovary deepCAGE and oocyte PAIso-seq data across transcript lengths from the 5' ends to the 3' ends and the flanking 2Kb regions. Each transcript is scaled to the same size and strand orientation. (H) Heatmap illustrating Pearson correlation coefficient of isoform-level expression in human (upper) and mouse (lower) oocytes which passed both TGS and NGS quality control (human: n = 20; mouse: n = 26). (I) Isoform diversity in genes related to meiotic cell cycle and spindle organization in human embryonic stem cell (ESC) and granulosa cell (GC).

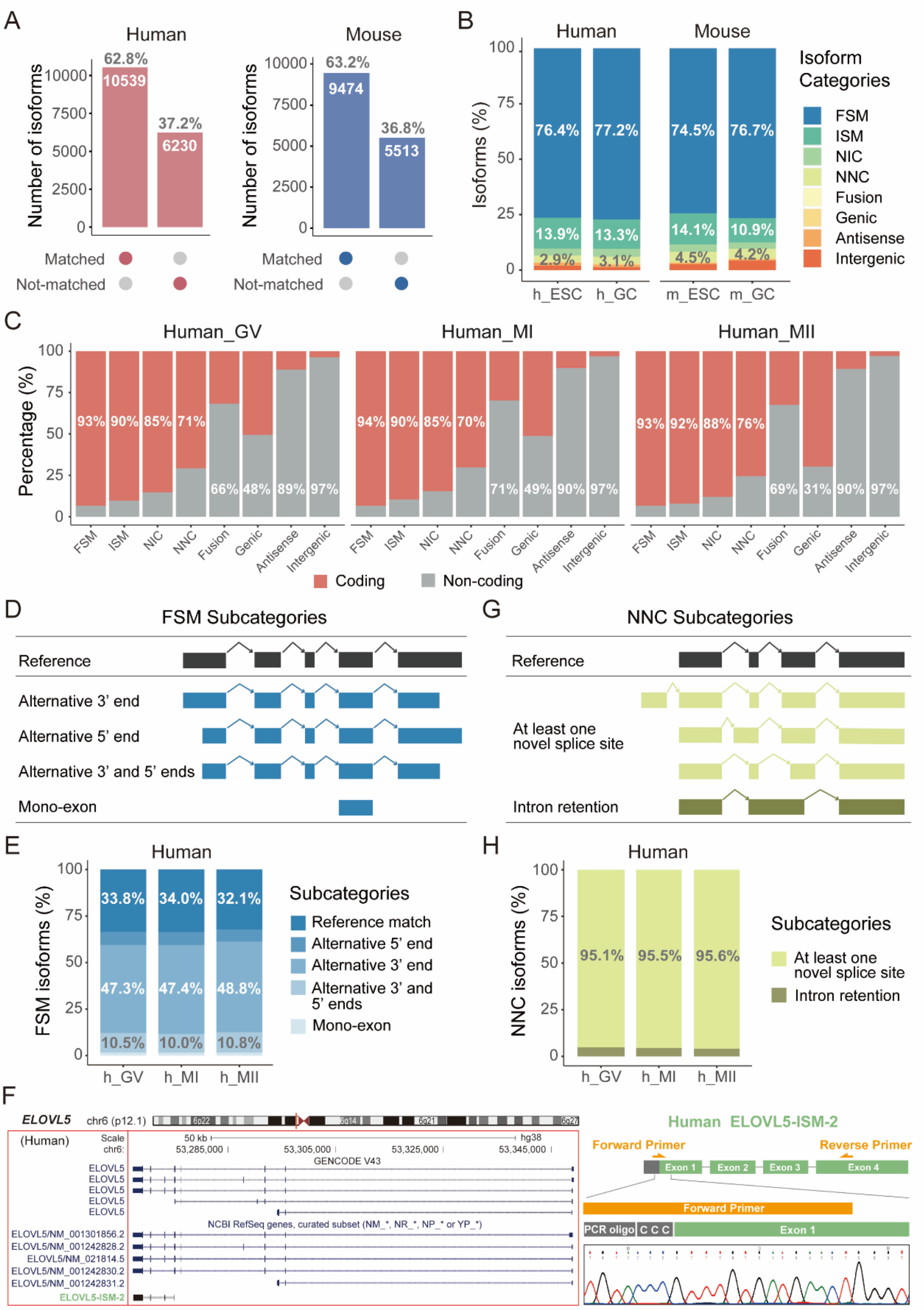


**Supplementary Figure 2. Isoform categories and subcategories of different isoforms in human oocytes.** (A) The comparison of isoforms detected in this study with previous transcriptomic data and GENCODE references during oocyte maturation. The number and proportion of matched and not-matched isoforms were labeled in the plot. (B) Percentages of different isoforms categories of ESC and GC in human and mouse. (C) Percentages of coding and non-coding isoforms vary across different isoform categories and human oocyte maturation stages. (D) Schematic diagram of subcategories of FSM. (E) Percentages of subcategories of FSM in different maturation stages of human oocytes. (F) The location in the genome and the isoform structure of *ELOVL5* are shown in the left. The schematic diagram of primer design and the junction between PCR oligo and transcript are shown in the right. The orange arrows showed the position of primers used for validation. (G) Schematic diagram of subcategories of NNC. (H) Percentages of subcategories of NNC in different maturation stages of human oocytes.

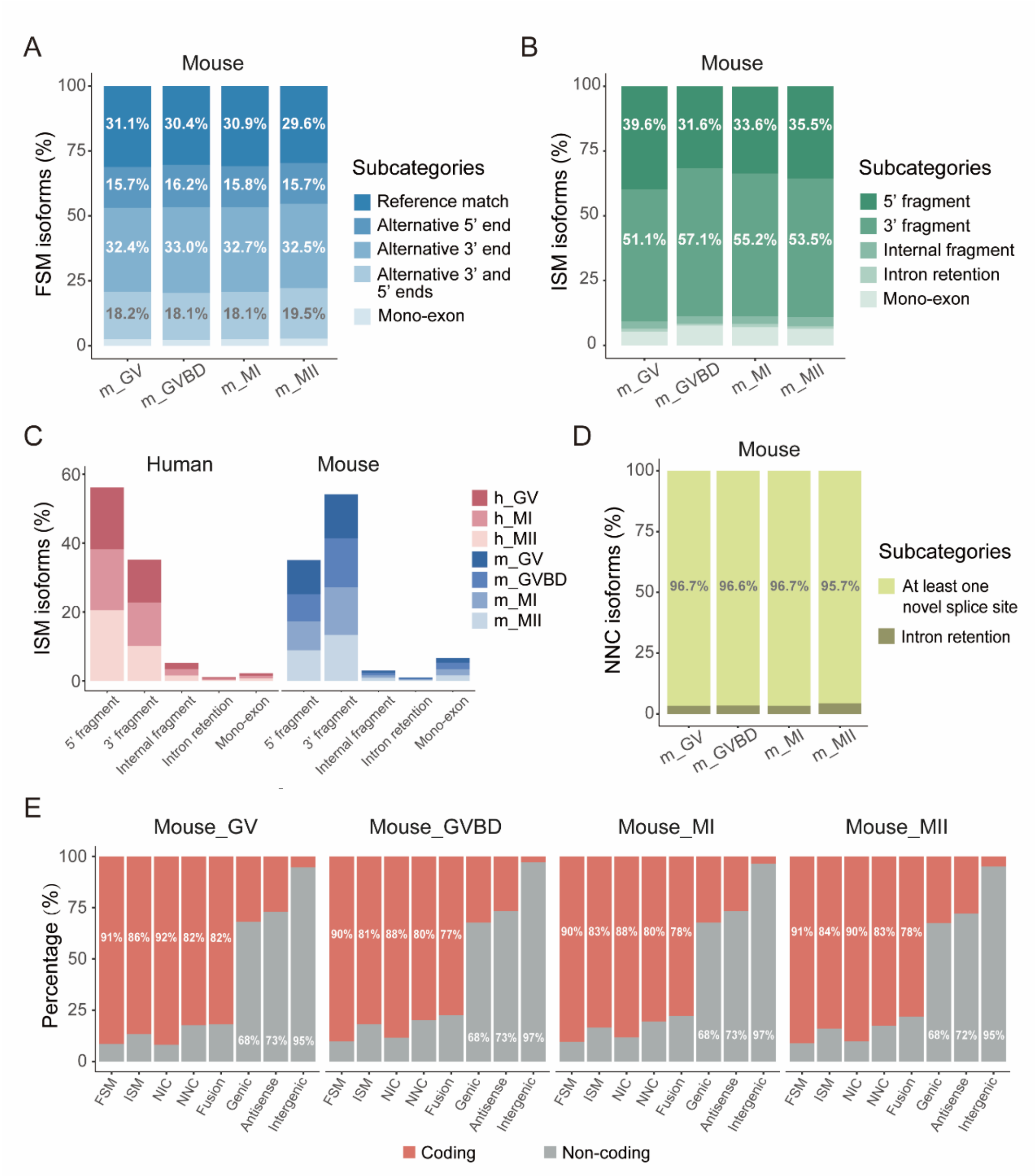


**Supplementary Figure 3. Subcategories of different isoforms in mouse oocytes.** (A) Percentages of subcategories of FSM in different maturation stages of mouse oocytes. (B) Percentages of subcategories of ISM in different maturation stages of mouse oocytes. (C) Fractions for each ISM subcategories in three human oocyte maturation stages (left) and four mouse oocyte maturation stages (right). (D) Percentages of subcategories of NNC in different maturation stages of mouse oocytes. (E) Percentages of coding and non-coding isoforms vary across different isoform categories and mouse oocyte maturation stages.

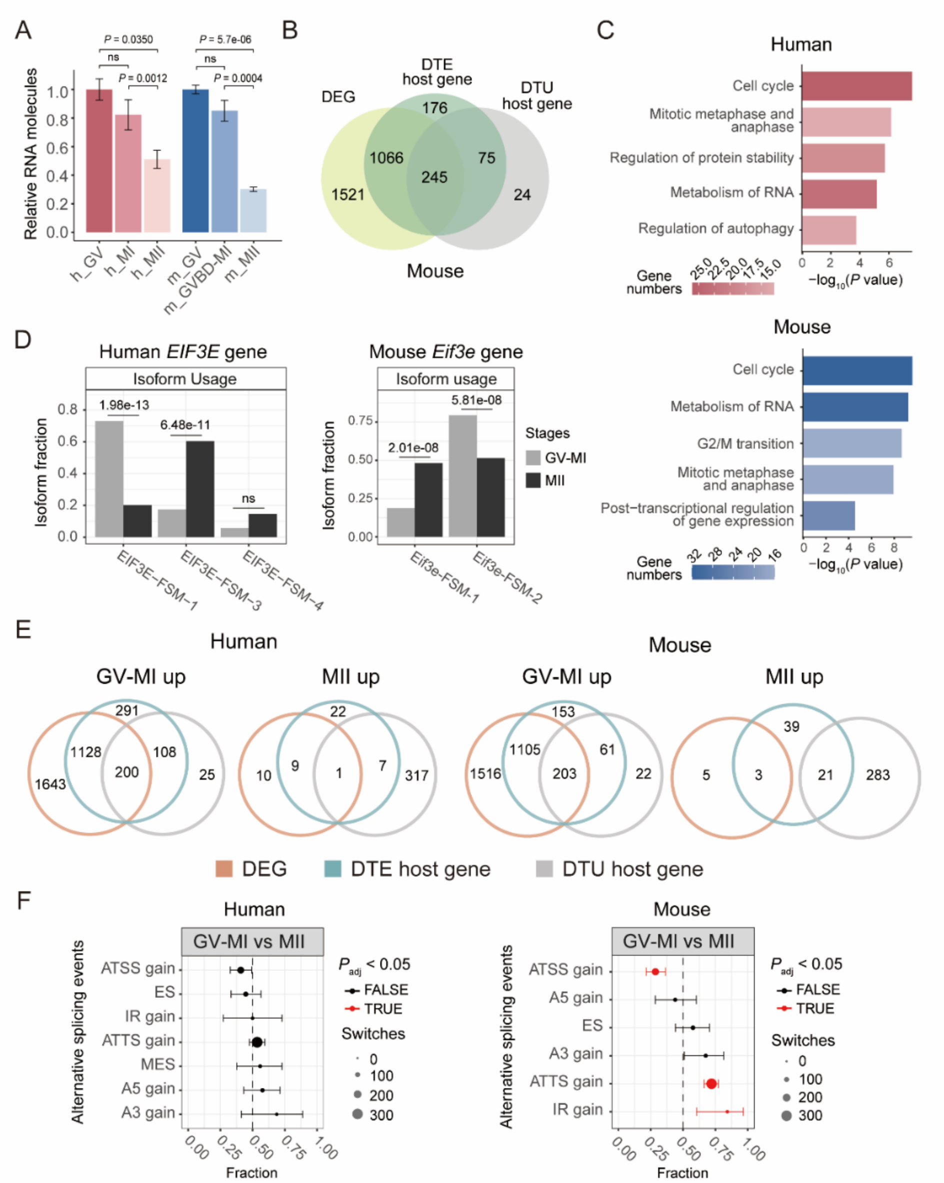


**Supplementary Figure 4. Comparative analysis of DTU events in human and mouse oocyte maturation.** (A) Bar plot showing maternal mRNA degradation during oocyte maturation in human and mouse. *P* values between different stages were obtained from Wilcoxon test (two-tailed unpaired; FDR-corrected). (B) Venn diagram illustrating the overlap of genes with significant DEG, DTE and DTU during mouse oocyte maturation. (C) Pathway enrichments for genes with DTU events during oocyte maturation in human (upper) and mouse (lower). (D) Bar plot showing isoform fraction of *EIF3E*/*Eif3e*, an example gene shared by both human and mouse exhibiting isoform fraction changes between the GV-MI and MII stages. Statistical significance of isoform fraction changes was evaluated with DEXSeq using a Likelihood Ratio Test based on a generalized linear model with a Negative Binomial distribution. *P* values were adjusted for multiple testing using the Benjamini-Hochberg method. Isoforms with IF values > 0.05 in at least one stage are plotted. (E) Venn diagram showing the overlap of upregulated genes identified as DEG, DTE, and DTU across different stages in human and mouse oocytes. (F) Splicing enrichment of isoform switches between GV-MI and MII stages in human (left) and mouse (right). The *x*-axis shows the fraction of events for each AS type. Significantly enriched splicing types are highlighted in red, with 95% confidence intervals, and the dot-size indicates the number of isoforms with certain type of change. *P* values were calculated using two-sided chi-square test with false discovery rate (FDR) corrected. DEG, differentially expressed gene; DTE, differential transcript expression; DTU, differential transcript usage; A5, alternative 5' donor site; A3, alternative 3' acceptor site; ATSS, alternative transcription start site; ATTS, alternative transcription termination site; ES, exon skipping; MES, multiple exon skipping; IR, intron retention.

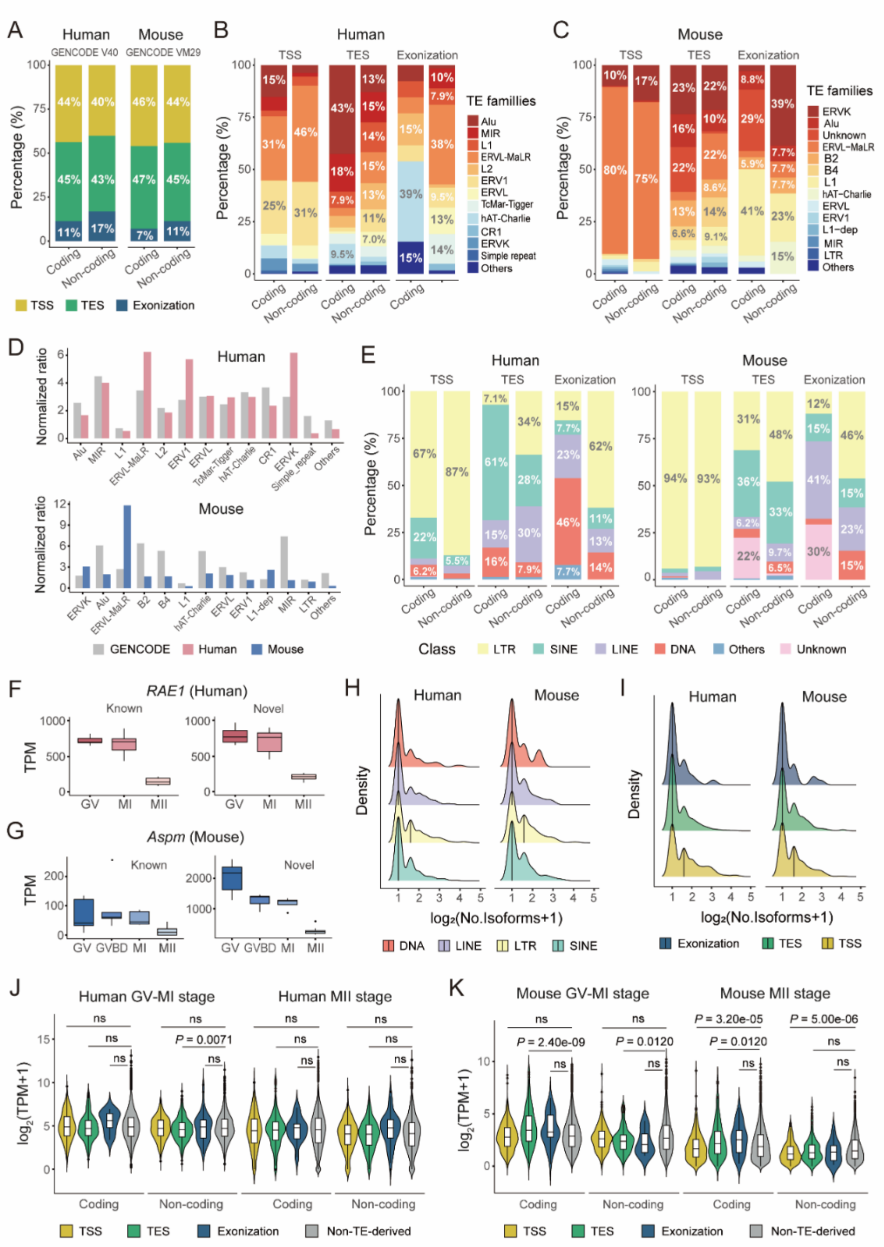


**Supplementary Figure 5. Comprehensive comparison of TE-derived and non-TE-derived isoforms among genome, GENCODE and oocyte assemblies.** (A) Percentages of different TE-derived isoform types in GENCODE V40 and GENCODE VM29 references (n = 47,422 in human; n = 19,914 in mouse), separated by coding probability. (B,C) Ratios of coding and non-coding isoforms derived by different TE families in TSS, TES, and exonization types in human (B) and mouse (C) oocytes. (D) The comparison of TE family members incorporated into oocyte transcripts assembled in this study and the GENCODE transcripts, normalized by their genomic distribution. (E) Ratios of coding and non-coding isoforms derived by different TE classes in TSS, TES and exonization types in human and mouse oocytes. (F) The expression levels of known and LTR-derived novel isoforms of *RAE1* during human oocyte maturation. (G) The expression levels of known and LTR-derived novel isoforms of *Aspm* during mouse oocyte maturation. (H) Density plot showing the number of transcripts that individual TE from different classes can be incorporated into in human and mouse oocytes. (I) Density plot showing the number of three types of TE-derived transcripts (TSS, TES and exonization) that individual TE can be incorporated into in human and mouse oocytes. (J,K) Comparison of the expression of non-TE-derived isoforms and TE-derived isoforms across various types in human (J) and mouse (K) oocytes, separated into coding and non-coding transcripts. The Wilcoxon test was used for significance testing, and $P$ values were corrected for multiple comparisons using the Benjamini-

Hochberg FDR method.

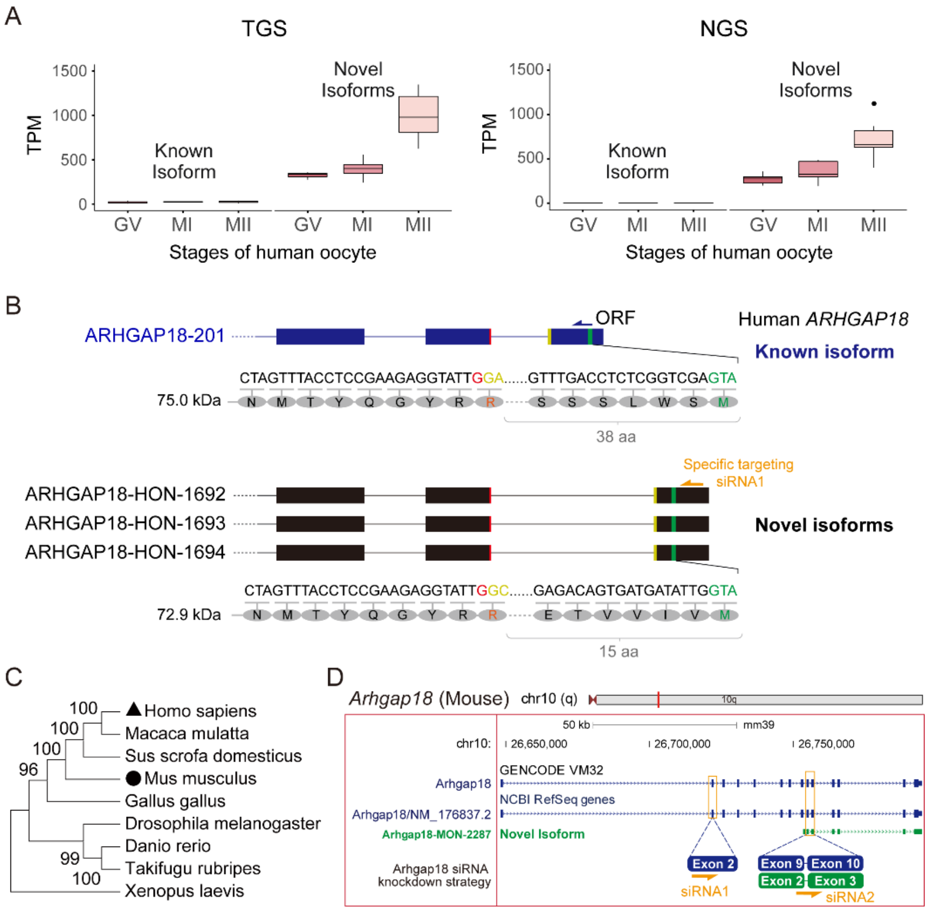


**Supplementary Figure 6. Additional information of the *ARHGAP18*/*Arhgap18* gene.** (A) Box plot showing the summed expression level of known and novel isoforms of *ARHGAP18* during oocyte maturation by TGS and NGS. (B) The different amino acid sequences at the N terminal of the protein coded by known and novel isoforms of *ARHGAP18*. The predicted protein weight coded by the novel isoform is 72.9 kDa. ORF, open reading frames; aa, amino acid. (C) Phylogenetic tree represents the evolutionary relationships of ARHGAP18 protein among various species. (D) Diagram illustrates the knockdown strategy of *Arhgap18* in mouse oocyte. The siRNA sequences are provided in the method section.
